# Transcranial magnetic vs intracranial electric stimulation: a direct comparison of their effects via scalp EEG recordings

**DOI:** 10.1101/2025.05.21.654985

**Authors:** Renzo Comolatti, Gabriel Hassan, Ezequiel Mikulan, Simone Russo, Michele Colombo, Elisabetta Litterio, Giulia Furregoni, Sasha D’Ambrosio, Matteo Fecchio, Sara Parmigiani, Ivana Sartori, Silvia Casarotto, Andrea Pigorini, Marcello Massimini

## Abstract

**Background:** Single-pulse Transcranial Magnetic Stimulation (TMS) and Intracranial Electrical Stimulation (IES) are widely used to probe cortical excitability and connectivity, but their electrophysiological effects have never been compared.

**Objective:** This study aims to fill this gap by using high-density scalp electroencephalogram (hd-EEG) as a common read-out to compare human brain responses to TMS and IES.

**Methods:** The dataset includes TMS-evoked potentials (TEPs) acquired from healthy subjects (n=22) and IES-evoked potentials (IEPs) recorded from drug-resistant epileptic patients (n=31) during wakefulness. In a subset of subjects TEPs (n=12) and IEPs (n=13) were also recorded during NREM sleep. Amplitude, spectral, and spatiotemporal features of TMS and IES responses, as well as their estimated electrical fields, were compared.

**Results:** We observed marked differences between TMS and IES responses. During wakefulness, IEPs are considerably larger, slower and associated with a suppression of cortical activity, whereas TEPs are characterized by multiple waves of recurrent activation. These differences are attenuated during NREM, during which both TMS and IES elicit large EEG responses associated with a prominent suppression of cortical activity. At the global level, the spatiotemporal complexity of the responses to both TMS and IES decreases consistently following the transition from wakefulness to NREM sleep.

**Conclusion:** Despite the limitations due to different subject populations (healthy vs pathological), our findings provide a first reference to parallel non-invasive and invasive brain stimulation and to interpret their differential effects. They also offer important insight on how cortical responsiveness is shaped by inhibition and adaptation mechanisms depending on input parameters and brain states.

**Highlights:**

- IES evokes higher signal-to-noise EEG responses than TMS.
- IES responses are larger, slower, and lead to cortical suppression.
- TMS elicits faster recurrent waves, with no suppression during wakefulness.
- Both TMS and IES converge to similar patterns during NREM sleep.

## Introduction

Single-pulse cortical stimulation in combination with electrophysiological recordings offers a unique window to explore the input-output properties of cortical neurons and their large-scale interactions from a causal perspective. This perturb-and-measure approach has been widely employed to investigate cortical plasticity, excitability and connectivity. In humans, single-pulse direct cortical perturbations have been implemented with non-invasive methods—as Transcranial Magnetic Stimulation (TMS), delivering magnetic pulses through the scalp (1,2)—and with invasive methods—as Intracranial Electrical Stimulation (IES), delivering electric pulses through surgically implanted electrodes. Both TMS and IES have been employed to probe cortical excitability (3–7), effective connectivity (8–11) and to map cortical (12–15) and subcortical (16) networks across different physiological and pathological conditions. Both techniques have also been employed to study altered states of consciousness such as NREM sleep (17–19), anesthesia (20–22), severe brain injury (23–25), epilepsy (26,27).

While TMS and IES share the fundamental characteristic of directly activating cortical neurons, they rely on different biophysical principles that may differentially shape the intensity and spatial extent of the stimulating field, leading to potentially divergent responses. Moreover, while most TMS studies have been performed in healthy subjects, IES is typically delivered based on clinical needs in patients with epilepsy. Directly comparing the electrophysiological effects of TMS and IES is thus key to interpret the current literature, align non-invasive and invasive approaches, and design future experiments. Yet, such a direct comparison has been so far hindered by the lack of a comparable read-out, as the cortical effects of TMS and IES have been previously recorded at different levels: scalp EEG in the first case (17,28) and intra-cranial EEG in the second case (11,13).

In the present study, we directly compare for the first time the amplitude, spectral and spatio-temporal features of TMS-evoked potentials (TEPs) and IES-evoked potentials (IEPs) at the common scale of high-density scalp EEG. Given established changes in cortical excitability across brain states, especially during NREM sleep, the study also examines how responses to stimulation evolve with changes in vigilance, offering mechanistic insight into state-dependent cortical dynamics. To this aim, recordings were performed during wakefulness and NREM sleep, while TMS and IES were delivered with stimulation parameters used in typical research and clinical protocols (11,23,28,29).

## Material and Methods

### Participants, data acquisition and preprocessing

Both IES-EEG and TMS-EEG studies were authorized by the local Ethical Committee (IES-EEG Prot.n. 348-24062020, Niguarda Hospital, Milan, Italy; TMS-EEG Prot.n. 609/07/27/05/AP, Comitato Etico Milano Area 1), and all participants provided written informed consent. Age-matching analysis confirmed no significant difference between the TMS and IES groups (see Fig. S1).

#### TMS-EEG

The TMS-EEG dataset included in the present study comprises 90 sessions collected from 22 healthy awake subjects (age = 31.4 ± 10.3; 13 F). In addition, 12 paired sessions were collected from 12 participants (age = 32.8 ± 8.3; 6 F) during NREM sleep, stage N3 (30), 10 of which previously published (23,31). These sessions were included based on the following criteria: (i) stimulation was delivered during a stable N3 sleep stage; (ii) a sufficient number of trials (>200) were registered to ensure a high signal-to-noise ratio, and (iii) a paired wakefulness session was also recorded. Targeted sites included Premotor (BA06), Parietal (BA07) and Occipital (BA19) cortices (see Table S1 for demographic and stimulation details of each subject). The data were recorded using TMS-compatible 64-channel amplifiers (Brain Products GmbH, 94 sessions, 5000 Hz sampling rate; Nexstim Ltd., 20 sessions, 1450 Hz sampling rate).

Each TMS session consisted of a minimum of 200 pulses (230±27) per area per subject administered with an inter-stimulus interval randomly jittered between 2 and 2.3 s. Biphasic pulses lasting 230 μs were delivered using a focal figure-of-eight coil at estimated E-field intensities of approximately 120 V/m (Fig. 1A).

**Fig. 1.**
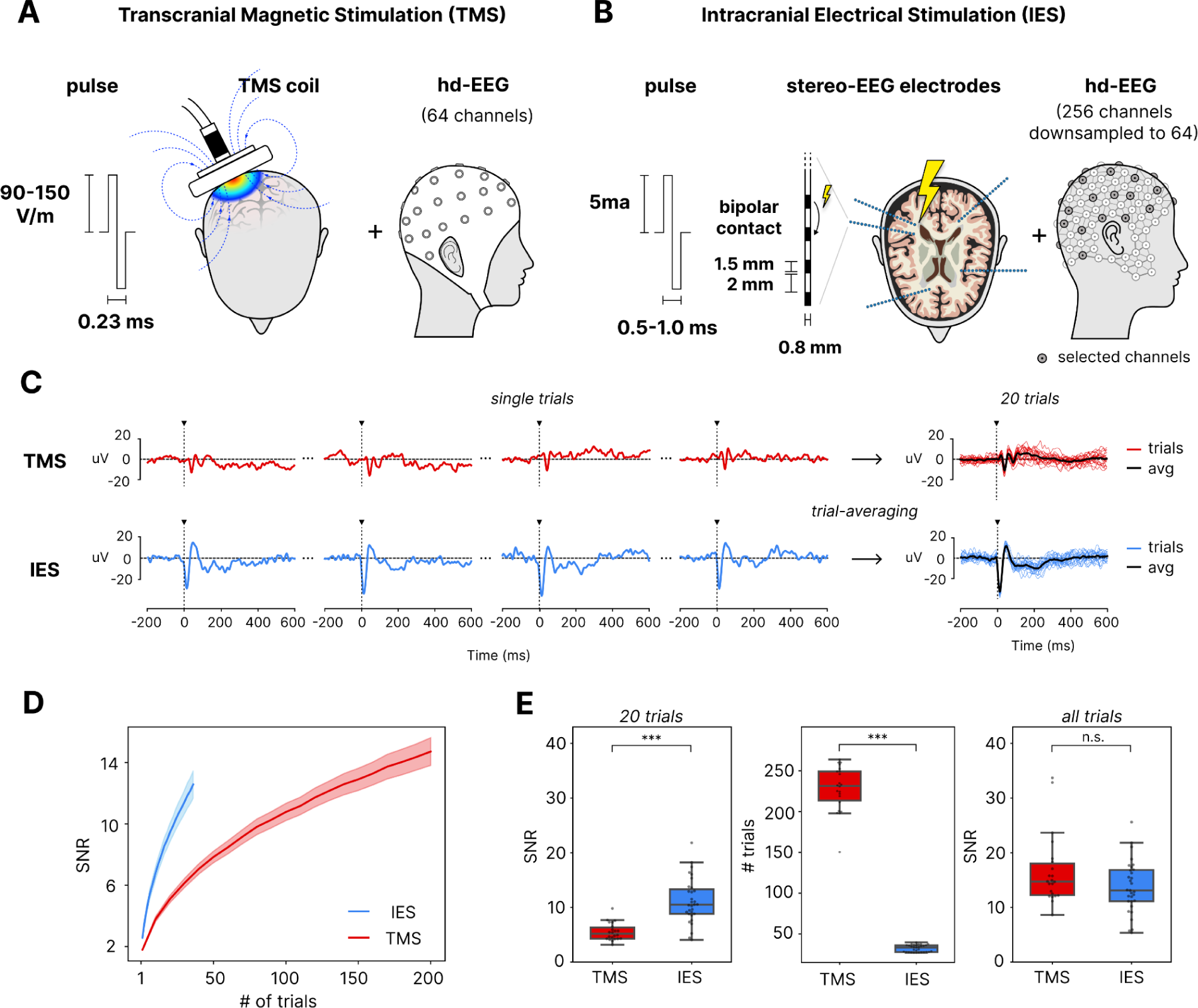
Experimental setup of Transcranial Magnetic Stimulation (TMS) and Intracranial Electrical Stimulation (IES) using hd-EEG and quantification of signal-to-noise ratio. (A) Schematic of the TMS-EEG procedure performed in healthy subjects depicting the biphasic pulse delivered with the TMS coil to the scalp, resulting in a magnetic field intensity between 90-150 V/m over a duration of 0.23 ms, and the recording setup with a 64-channel hd-EEG. (B) Illustration of the IES-EEG setup used on epileptic patients undergoing presurgical evaluation comprising the intracranially implanted stereotactic electrodes (SEEG) which deliver biphasic pulse of 0.5-1.0 ms duration through bipolar contacts (separated by 2mm) at constant intensity of 5mA, alongside the simultaneous recording setup with a 256-channel hd-EEG. (C) Scalp EEG traces from single trials of TMS (top) and IES (bottom) for the strongest responding channel, showing time-locked potentials elicited by the single-pulses (black triangle). To the right, the evoked average (black trace) of 20 single trials (colored traces) depicting the build up of signal-to-noise (SNR) through trial-averaging. (D) Relationship between SNR and the number of trials utilized for trial-averaging, obtained by bootstrapping procedure. (E) Boxplots depicting SNR comparisons for TMS and IES after 20 trials (left), the total number of trials in each recording (middle), and SNR from all trials (right). Statistical significance is denoted by asterisks (n.s.=p>0.05, * = p<0.05, ** = p<0.01, *** = p<0.001)

TMS-EEG data were processed similarly to (31). The stimulation artifact was first removed and hd-EEG data was high-pass filtered at 0.5 Hz, splitted into epochs and bad trials and channels were rejected by visual inspection. Epochs were re-referenced to the average reference and baseline corrected. After Independent Component Analysis (ICA) was applied to remove EMG and EOG activity, the signal was low-pass filtered at 45 Hz and down-sampled to 1000 Hz. See Supplementary Materials for further details.

#### IES-EEG

The IES-EEG dataset was obtained from patients undergoing intracranial monitoring for pre-surgical evaluation of drug-resistant epilepsy (32). A total of 231 sessions were acquired from 31 subjects (mean age = 31, 18 F) during wakefulness (data available at https://osf.io/wsgzp/ (6); see Table S2 for demographic and clinical details of each patient). In a subset of 13 patients (mean age = 28, 9F), 54 paired sessions were also acquired during NREM sleep, stage N3 (30), as in (18). Stimulation locations were spread across the entire cortical mantle (see (6)) including the areas targeted by TMS. Each session consisted of circa thirty bipolar biphasic pulses (32±4.2) of 500 and 1000 μs duration delivered at intervals ranging from 1 to 5 s. Pulses were delivered through stereotactically implanted (SEEG) intracerebral electrodes at 5 mA intensity between adjacent contact pairs. Recordings were simultaneously conducted using 256 channels high-density scalp EEG (1000 Hz sampling rate) and downsampled to 61 channels (33), equivalent to the ones used for TMS-EEG data collection (Fig. 1B).

IES-EEG data were processed similarly to TMS-EEG data as in (6). First, channels and trials contaminated by noise, muscle activity or spontaneous interictal epileptic discharges were rejected using a semi-automatic procedure, manually verified by an expert electrophysiologist. Next, the stimulation artifact was removed and data were band-pass filtered (0.5-45 Hz) and epoched. Finally, trials were re-referenced to the average and baseline corrected and ICA was applied to remove EOG activity. See Supplementary Materials for further details.

### Data Analysis

Signal-to-noise ratio (SNR) measures signal strength time-locked to stimulation with respect to the pre-stimulus background activity. SNR was calculated as the square root of the ratio of average power between the early response (0 to 80 ms) and the baseline (−300 to −5 ms), at the channel with the highest power in the initial 80 ms. The influence of the number of trials on the SNR was evaluated using a bootstrap method: for a given number of *k* trials, SNR was computed on 100 surrogate responses formed by randomly selecting *k* trials without replacement.

The Global Mean Field Power (GMFP) estimates the overall power evoked at each time point across all channels and corresponds to the spatial standard deviation of the signal (34). The GMFP was then baseline corrected to quantify the power of the evoked response exceeding the power in the spontaneous baseline activity. The Perturbational Complexity Index state-transition (PCI^ST^) was used to assess the spatiotemporal complexity of the evoked potentials (35). PCI^ST^ gauges the ability of thalamocortical circuits to engage in complex causal interactions, by jointly quantifying the spatial diversity and temporal differentiation of brain responses. PCI^ST^ was computed over the 0-600 ms response window with remaining parameters following (35). Code available at github.com/renzocom/PCIst. See Supplementary Materials for further details.

We evaluated the modulation of high-frequency (≥ 20Hz) EEG oscillations induced by the stimulation using the event-related spectral perturbation (ERSP) method (36) as in (31). We calculated the high-frequency power (HFp) for each channel as the average time course of significant deviations from baseline (bootstrap; α ≤ 0.01, 500 permutations) in instantaneous high-frequency power between 120 and 220 ms. From the distribution of HFp across channels we computed three metrics to gauge the suppression of high-frequency: (i) the *extent* of suppression was measured as the percentage of suppressing (HFp < 0) channels (%Ch HFsup); (ii) its maximal *intensity* as the minimum HFp value across all channels (max HFsup); and (iii) its *total* amount as the integral of HFp < 0 across channels normalized by the total number of channels (total HFsup). We applied all these high-frequency suppression measures above 20 Hz with the aim of gauging the “OFF-periods” as defined by (18) at the intracerebral level and by (31) at the TMS-EEG level. This approach builds on previous animal and human intracranial recordings, which have shown that the silent, hyperpolarized state characterizing the cortical OFF-periods during spontaneously occurring sleep slow oscillations is associated with a suppression of high-frequency power (>20 Hz) in the local field potential (LFP) (37–42) (see also Fig. S5).

### Statistical analysis

All metrics were computed on the individual evoked potentials. Then, metrics were averaged across stimulation sessions for each subject and then tested across groups using t-tests. Specifically, differences between TMS and IES, which involve different numbers of subjects, were assessed using Welch’s t-test. Conversely, comparisons between wakefulness and NREM sleep, within stimulation method, were conducted using a paired Student’s t-test, correcting for multiple comparisons using the False Discovery Rate (FDR) method. Data reported as mean±std. To ensure the robustness of our findings, we additionally confirmed the results using Linear Mixed Models (LMMs), as detailed in the Supplementary Material (see Supplementary Methods and Tables S3-13).

### Simulation of electric field

E-fields are instantaneous estimates of the electrical potential gradients induced in the brain tissue by the two stimulation devices—the magnetic coil in TMS and the bipolar contacts of the SEEG electrodes for IES. To ensure consistency in the comparison, the TMS and IES E-fields were computed with the finite element method (FEM) using a common realistic volume conductor model based on the MNI152 template (43), with conductivities set at 0.14 S/m for white matter and 0.33 S/m for gray matter (44).

Given the fundamental differences in how TMS and IES generate electric fields, we used SimNIBS 4.0 (45) for TMS and Lead-DBS 3.0 (46) for IES, as each is specifically designed for its respective stimulation modality. TMS E-field was simulated using a 70 mm figure-of-eight coil template (MagVenture MC-B70—the most similar to the coil used in the experiments), while IES E-field was modeled using a template of the electrode used in the clinical studies (Dixi Medical, as in Fig. 1B). Both employ FEM-based simulations, ensuring technically accurate and physiologically meaningful estimations.

Simulations were conducted at standard (120 V/m for TMS and 5mA for IES) and high intensities (160/Vm and 10mA, respectively). The TMS coil was placed over the crown of a cortical gyrus with the field oriented perpendicular to it to maximize the cortical efficacy (47). Intracranial bipolar contacts for IES stimulation were positioned in the gyrus aligned with the TMS coil’s normal projection. The E-field peak strength was defined as the 99.9% percentile of the field strength distribution to avoid potential outliers in the FEM simulation (47). The spatial decay of the E-field was calculated by finding, for every E-field threshold value, the farthest distance in 3D anatomical coordinates that was still above it.

## Results

We compared the effects of single-pulse TMS and IES by first analyzing scalp hd-EEG responses registered during wakefulness. For both TMS and IES, stimulation parameters followed the typical settings used to effectively elicit cortical responses in experimental and clinical settings within safe operational ranges (for example (23,28) for TMS, and (6,11,19) in the case of IES; see Methods).

### IES evokes EEG responses with higher signal-to-noise ratio than TMS

As displayed in Fig. 1C, both TMS and IES induced time-locked potentials that were visible from the scalp EEG at the single trial level and reproducible across trials. Once trial averaged, both TMS and IES responses yielded evoked potentials with high signal-to-noise ratio (SNR) (Fig. 1C, right). Nonetheless, TEPs were less prominent with respect to the pre-stimulus baseline than IEPs. The SNR of IEPs increased more rapidly as a function of the number of trials averaged than TEPs (Fig. 1D). After averaging twenty trials, IEPs showed a twofold difference in SNR with respect to TEPs (TMS=5.5±1.6, IES=11±4.01, p=1.7×10^-8^) (Fig. 1E, left). This difference was offset by the larger number of trials collected in the case of TMS (TMS=230±27, IES=32±4.2, p=1.1×10^-39^) (Fig. 1E, middle), resulting in comparable SNR (TMS=17±6.3, IES=14±4.7, p=0.09) that did not differ significantly once all trials were considered (Fig. 1E, right).

### IES evokes EEG responses that are larger than those evoked by TMS

At comparable SNR, trial-averaged evoked potentials revealed both commonalities as well as noticeable differences between TMS and IES. Fig. 2A depicts, on a common scale, the butterfly plots of individual trial-averaged TEPs and IEPs from different brain regions (occipital, parietal and premotor) of representative subjects, overlaid by the respective global mean field power (GMFP) (black traces). Both stimulation methods evoked high-amplitude and long lasting potentials, displaying a composite response with non-trivial spatiotemporal activation profiles, which varied across stimulation sites. Nonetheless, the global power of IEPs was strikingly larger than that of TEPs, as illustrated by the direct comparison between the grand averages of the GMFP evoked by the two stimulation modalities (Fig. 2B). Significant differences were found when considering the average GMFP across the whole time course (TMS=0.55±0.13 uV, IES=3.7±2.0 uV, p=1.3×10^-9^) (Fig. 2C, left), as well as when considering separately the peak amplitudes of the early (0-80 ms, TMS=3.0±0.6 uV, IES=8.5±2.8 uV, 3.6×10^-12^) (Fig. 2C, middle) and late (80-600 ms, TMS=1.6±0.44 uV, IES=9.8±5.7 uV, p=9.2×10^-9^) components of the GMFP (Fig. 2C, right). These differences in the GMFP between TMS and IES remained consistent across stimulated regions (see Fig. S3).

**Fig. 2.**
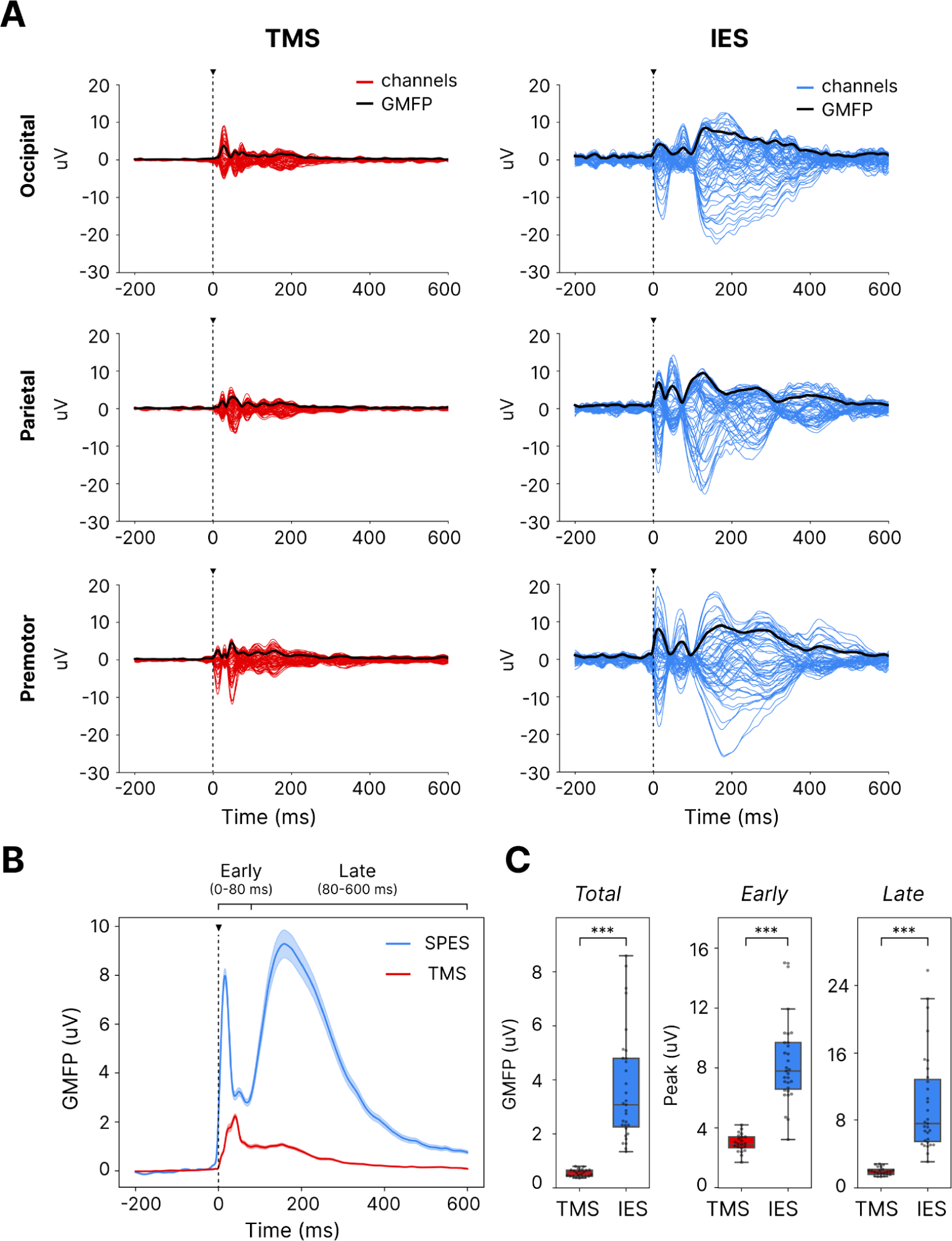
GMFP comparisons between TMS and IES Evoked Potentials. (A) Butterfly plots of individual TEPs and IEPs obtained by the stimulation of occipital, parietal, and premotor areas in representative subjects. The colored traces depict the evoked response across EEG channels on a common scale, with the GMFP overlaid in black, illustrating the power evoked globally across the EEG channels over time (see Fig. S2 for a zoomed in view of signals, auto-scaled and in a shorter time interval). (B) Grand averages of the baseline corrected GMFP time course for all TEPs (red) and IEPs (blue) and their respective standard errors (light shadow), illustrating the differences in evoked power over time, segmented into early (0-80 ms) and late (80-600 ms) intervals. (C) Boxplots compare the total GMFP across the full response interval (0-600 ms), and the peak GMFP amplitudes for the early (0-80 ms), and late (80-600 ms) intervals. Statistical significance is denoted by asterisks (n.s.=p>0.05, * = p<0.05, ** = p<0.01, *** = p<0.001).

### The estimated electric field is weak and widespread for TMS, strong and focal for IES

The common read-out of scalp EEG recordings revealed a substantial difference in the magnitude of the overall response evoked by TMS and IES. We next investigated this finding in light of the differences in the strength and spatial extent of the electric fields (E-fields) generated in the cortex by the two stimulation modalities (see Methods for details). Fig. 3A depicts the simulated E-field of a parietal cortex stimulation (BA07) at standard stimulation intensity for TMS (Fig. 3A, *top*) and IES (Fig. 3A, *bottom*) in a common scale.

**Fig. 3.**
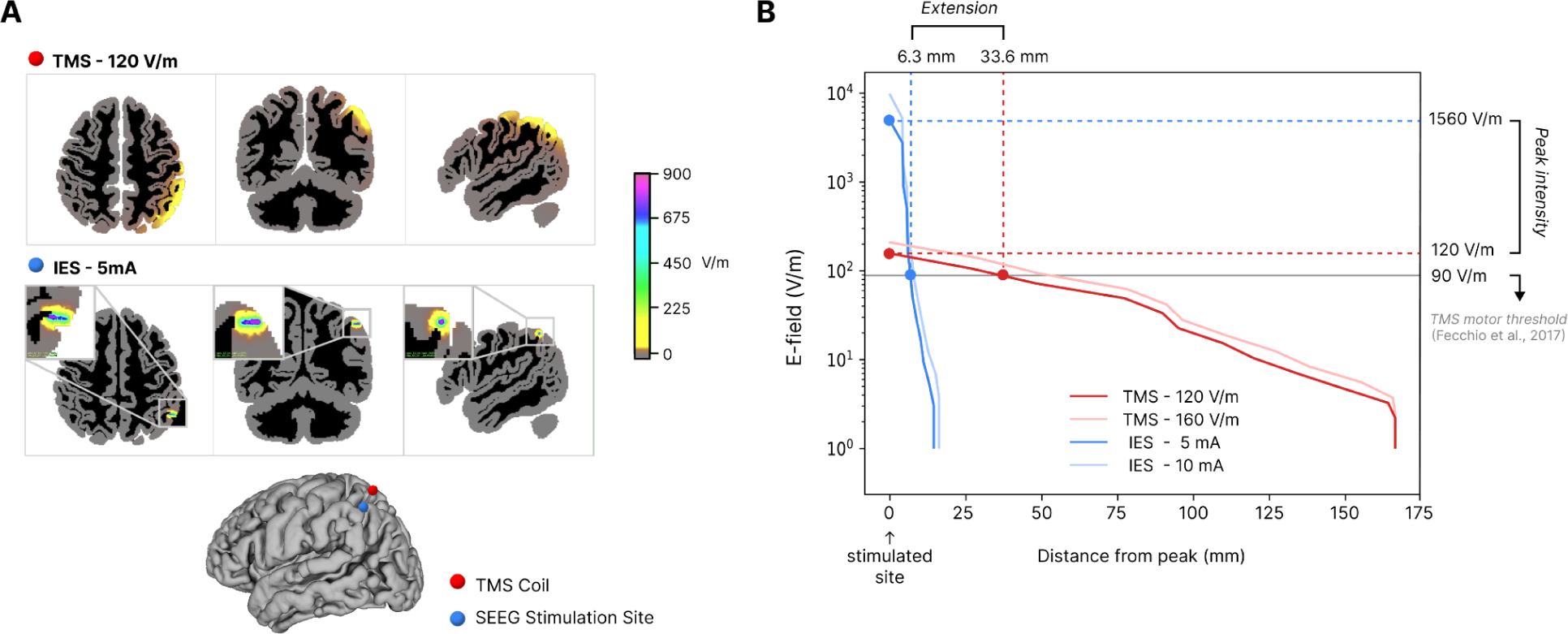
Simulated electric fields induced by TMS and IES. (A) Simulated E-fields induced during standard intensity stimulation to parietal cortex by TMS and IES on a MNI152 template. The upper panels show E-fields for IES at 5mA, highlighting the focused high amplitude area close to the stimulation bipolar contacts. The lower panels depict the more diffuse and weaker E-field for TMS at 120 V/m stimulation intensity. The placement of the TMS coil (red) and SEEG stimulation site (blue) are indicated on a 3D brain model (bottom). (B) Spatial decay of E-field with distance from the stimulated site for TMS (red) and IES (blue), for both standard and high stimulation intensity. The horizontal dashed lines indicate maximum E-field strength (intensity), and vertical dashed lines indicate the E-field strength at an operational threshold of 90 V/m (gray line) (extension).

The E-field pattern of the two stimulation differed substantially: the E-field computed for IES showed a high intensity profile concentrated at the stimulation site, whereas the E-field computed for TMS was weaker but more widespread. Specifically, the maximum E-field value of IES was more than an order of magnitude greater than that of TMS (peak of E-field, TMS=120 V/m, IES=1560 V/m). On the other hand, the E-field induced by TMS was significantly less focal, exhibiting a more gradual spatial decay (Fig. 3B). For example, at 90 V/m—the typical E-field strength of the resting motor threshold in TMS (48). These differences were preserved across stimulation intensities (Fig. 3B) and sites.

### The amplitude of IEPs and TEPs differs in wakefulness but becomes more similar in NREM sleep

Having explored the differences in the magnitude of the response and the different E-field patterns generated by IES and TMS, we moved to investigate the effects of changes between brain states. To this aim, we analyzed a cohort of paired TEPs and IEPs recorded during wakefulness and NREM sleep.

During wakefulness, TMS elicited faster EEG components with a richer spatiotemporal profile (Fig. 4A) as compared to IES, as indicated by the higher number of peaks in the GMFP (TMS=7.0±1.1, IES=5.9±0.70, p=4.2×10^-4^). Upon falling asleep, the EEG response to TMS became larger and slower, and thus more similar to the one triggered by IES (Fig. 4B). The average GMFP of both TEPs and IEPs was higher during NREM sleep than during wakefulness (TMS_wake_=0.67±0.19 uV, TMS_sleep_=3.8±2.9 uV, p=6.0×10^-3^; IES_wake_=4.4±3.3 uV, IES_sleep_=6.2±3.2 uV, p=3.7×10^-4^). The GMFP of TEPs during NREM sleep resembled the GMFP profile observed in IEPs in both wakefulness and NREM sleep, being characterized by an early peak and a slow sustained late activation. Accordingly, the total GMFP of TEPs recorded during NREM sleep was not significantly different from the GMFP of IEPs recorded during both wakefulness (p=0.65) and NREM sleep (p=0.077)—this was also the case for early and late GMFP peaks (see Fig. S4).

**Fig. 4.**
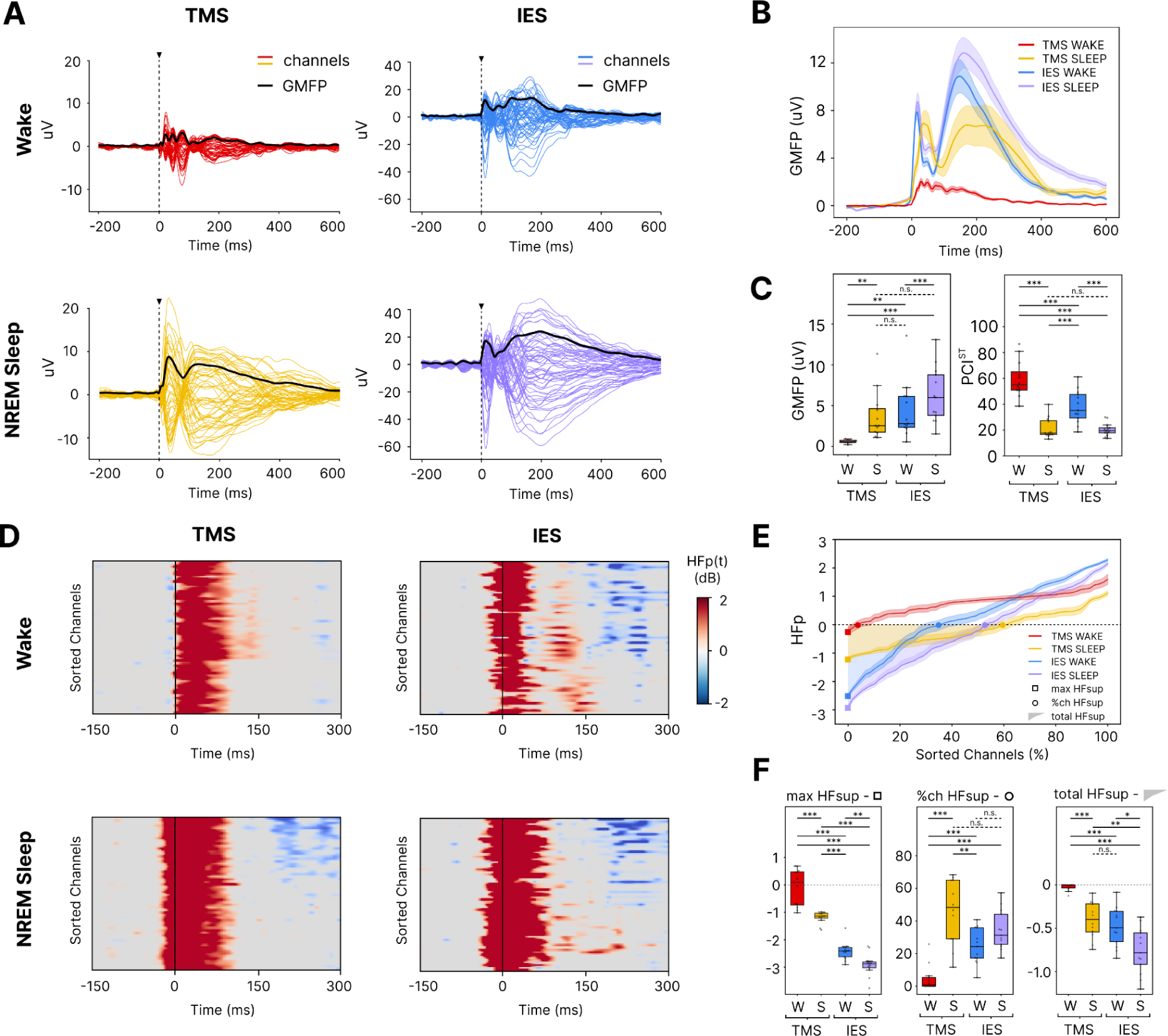
State-Dependent Changes in TMS and IES Evoked Responses. (A) Butterfly plots of representative TMS (left column) and IES (right column) evoked potentials during wakefulness (top) and NREM sleep (bottom). Each trace represents a single EEG channel response with the GMFP overlaid as a black line, highlighting the differences in waveform and complexity between states and stimulation types. (B) GMFP time course for TMS and IES during wakefulness and NREM sleep, with shaded areas indicating standard deviation. These illustrate changes in GMFP amplitude across brain states. (C) Boxplots of GMFP and PCI^ST^, quantifying the evoked potential’s amplitude and complexity, respectively, for TMS and IES during wakefulness (W) and NREM sleep (S). (D) High-frequency power (>20Hz) time courses across EEG channels during wakefulness and NREM sleep for TMS and IES, with blue indicating suppression (negative dB values) and red indicating an increase (positive dB values). Channels are sorted by their respective high-frequency power averaged across 120-220 ms (HFp). (E) Distributions of HFp sorted across channels for TMS and IES during wakefulness and NREM sleep (traces depict average and standard errors). Highlighted are the extent (circle, %Ch HFsup), maximum degree (square, Max HFsup) and overall (shaded area below zero, Total HFsup) suppression of high-frequency for each condition. (F) Boxplots show the metrics derived from the HFp distribution across channels: the proportion of channels showing suppression (%Ch HFsup), the maximal suppression across all channels (max HFsup), and the integral of HFp suppression across channels normalized by total number of channels (total HFsup), across conditions and stimulation types. These metrics gauge the extent, intensity, and overall suppression of HFp across channels, comparing TMS and IES during wakefulness and NREM sleep. Statistical significance is denoted by asterisks (n.s.=p>0.05, * = p<0.05, ** = p<0.01, *** = p<0.001).

### The spatiotemporal complexity of TEPs and IEPs differs but shows consistent changes across brain states

We then assessed the richness of the spatial and temporal profile of the evoked responses by computing the Perturbational Complexity Index (PCI^ST^) (35,49). During wakefulness, PCI^ST^ values were systematically higher for TEPs than IEPs (p=8.9×10^-4^) reflecting the richer spatiotemporal profile observed in the TEPs and the slower and more stereotypical features of IEPs. In line with previous findings (35), however, PCI^ST^ revealed state-dependent changes in complexity for both TMS and IES, being consistently higher in wakefulness than in NREM sleep (TMS_wake_=59±14, TMS_sleep_=21±7.7, p=1.6×10^-5^; IES_wake_=38±12, IES_sleep_=20±4.9, p=8.3×10^-5^).

Notably, the PCI^ST^ computed on IEPs during wakefulness remained consistently above the values obtained during NREM sleep in both IES and TMS (p=4.1×10^-3^). Lastly, consistent with the observed convergence of TEP and IEP’s waveform, the difference in PCI^ST^ values between TEPs and IEPs was non-significant during NREM sleep (p=0.68).

### Unlike TMS, IES induces suppression of high-frequency activity also during wakefulness

Last, we analyzed the modulation of high-frequency power (>20Hz, HFp) in the responses as a proxy for the suppression or increase of neuronal activity induced by TMS and IES (37,40). We used a frequency range of 20–45 Hz rather than >50 Hz, as is typically employed in classic intracerebral recordings (19,50), because in scalp EEG these frequencies better capture the dynamics of neuronal silence (i.e., OFF-periods) that we aim to observe (see for example (31,48,51)). Notably, these frequencies have also been used at the intracerebral level to gauge the extent of neuronal silencing (18,37). Still, a direct comparison between the two frequency ranges at the SEEG level did not reveal any substantial difference (Fig. S5A). Along the same line, within the 20–45 Hz range we observe a strong similarity between the high-frequency suppression seen at the scalp electrode with the largest amplitude and the underlying SEEG electrode in closest proximity (Fig. S5B). Fig. 4D displays the high-frequency time courses of each channel sorted by their respective HFp, for representative signals of each condition (see panel A). In the case of TMS, an initial broadband activation is followed by a suppression of HFp only in NREM sleep but not in wakefulness, as in (31,48). Conversely, the IES-induced initial activation was invariably followed by a clear-cut suppression of HFp both in wakefulness and NREM sleep (18). The distribution of HFp across EEG sensors depicted in Fig. 4E shows that, while HFp remained positive in almost all channels in the case of TMS during wakefulness (>95%), IES induced suppression of high-frequency in about 25% of channels during wakefulness. During NREM, instead, both TMS and IES induced a suppression of high-frequency in more than a third of all channels.

The extension (%Ch HFsup), intensity (maxCh HFsup) and overall amount (total HFsup) of high-frequency suppression was in general significantly stronger during NREM sleep compared to wakefulness for both TMS and IES (%Ch HFsup, TMS p=5.5×10^-5^, IES p=0.072; max HFsup, TMS p=4.3×10^-4^, IES p=9.5×10^-3^; total HFsup, TMS p=1.1×10^-4^, IES p=2.3×10^-2^) (Fig. 4F). During wakefulness, TMS responses displayed little to no suppression of high-frequency (total HFsup=-0.024±0.038 dB), both in terms of its extension across channels (%Ch HFsup=4.7±7.6 %) and maximal intensity (max HFsup=-0.097±0.65 dB). In contrast, IES responses exhibited a significant amount of high-frequency suppression during wakefulness, comparable to the total high-frequency suppression observed in NREM sleep for TMS (total HFsup, IES_wake_=-0.49±0.24 dB, TMS_sleep_=-0.41±0.2 dB, p=0.35). Taken together, these results suggest that while IES induces significant suppression of high-frequencies in both wakefulness and NREM sleep, TMS does so only during NREM sleep.

## Discussion

By maintaining scalp EEG as the common read-out for both TMS and IES, we found major differences in the magnitude and nature of their cortical effects. Seen from the scalp, IES elicits responses that are slower and up to an order of magnitude larger than TMS. In spite of major differences in amplitude, waveshape and spectral features, the global complexity of both TEPs and IEPs decreases from wakefulness to sleep. These results provide important insight on the mechanisms of cortical responses to direct perturbations across different stimulation parameters and brain states and draw a first parallel between non-invasive and invasive stimulation approaches.

### The interaction between stimulation strength and activity dependent mechanism shapes cortical responses

During wakefulness, TMS evoked multiple waves of recurrent activation, whereas IES triggered an initial activation (corresponding to the N1 component recorded with LFP, see (6,13)) that was rapidly followed by a large negative wave (corresponding to the N2 component recorded with LFP, see (6,13)) and by a concurrent suppression of high-frequency activity. As demonstrated by intracranial and extracranial studies in animals and humans, large EEG negative waves associated with high-frequency suppression correspond to the occurrence of a silent period (OFF-period) in cortical neurons (37,40). This tendency of cortical neurons to fall into a silent OFF-period after an initial activation reflects activity-dependent mechanisms such as adaptation Na+/Ca++ dependent K+ currents (52,53) and/or active inhibition (54). The differential effects of IES and TMS during wakefulness can be explained by the distinct E-field profiles generated by each stimulation technique (Fig. 3) and by their interactions with activity-dependent mechanisms (Fig. 5). The IES electric field is up to five times more focal than that induced by TMS and, at the typical stimulation parameters used in research and clinical settings, over ten times more intense. A parsimonious explanation is that the intense and highly-focused E-field (55,56) induced by IES strongly recruits activity-dependent mechanisms resulting in an OFF-period during wakefulness (57,58). The convergence between IEPs and TEPs observed during NREM sleep, can be explained by changes in neuromodulation leading to increase in both Na+/Ca++ dependent K+ currents (53,59) and inhibition (60,61) in cortical circuits. Such enhancement of activity-dependent mechanisms reduces the range of local activation levels that cortical circuits can withstand. This is consistent with the observation that during sleep even the relatively weaker TMS input is inescapably followed by an OFF-period.

**Fig. 5.**
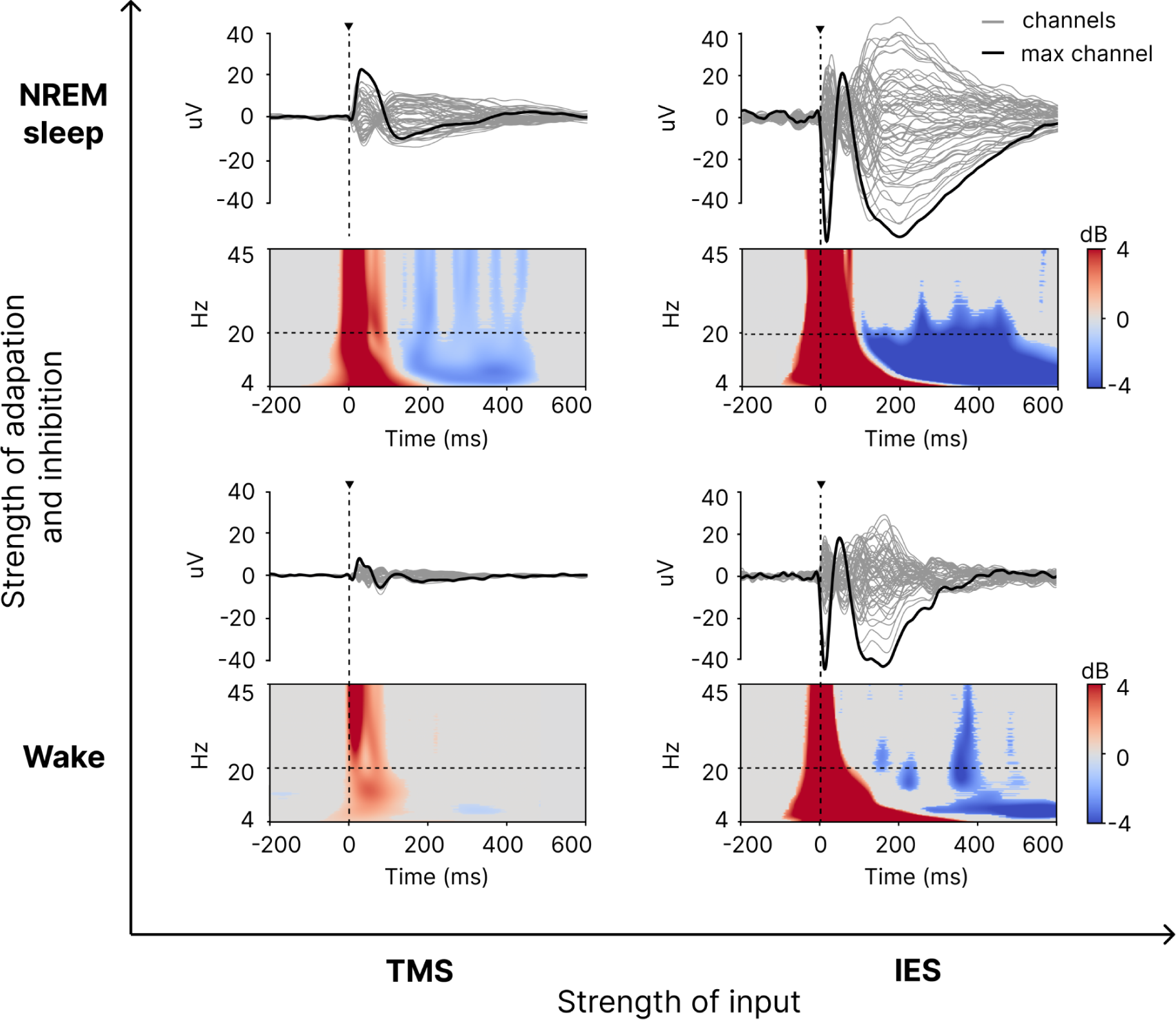
Input- and state-dependent modulation of TMS and IES evoked responses in wakefulness and NREM sleep. The x-axis represents the strength of stimulation input, increasing from left to right, with TMS (left column) providing a weaker, more diffuse stimulation compared to IES (right column), which delivers a stronger and more focal input. The y-axis represents the strength of adaptation (e.g. Na+/Ca++ dependent K+ currents) and inhibition mechanisms engaged in response to stimulation, which is lower in wakefulness (bottom row) and higher during NREM sleep (top row). In each quadrant, the upper panel depicts butterfly plots of representative evoked potentials, with all EEG channels (gray traces) and the channel with the higher amplitude (black trace). The black triangle indicates stimulus onset. The lower panel depicts the time-frequency ERSP plot of the strongest responding channel. Statistically significant increases of power compared to baseline are depicted in red, while blue represents significant power decreases. Absence of any significant activation is shown in gray. The dashed horizontal line indicates the 20 Hz frequency bin above which modulation of high-frequency power (HFp) is assessed (see Methods).

This aligns with computer simulations pointing to the importance of activity-dependent adaptation mechanisms in shaping cortical responsiveness (52) and with the general notion that cortical circuits’ dynamic range and information capacity decreases during NREM sleep (62,63). Similarly, in conditions like stroke and traumatic brain injury, where adaptation and inhibition increase, TMS can induce sleep-like cortical responses during wakefulness, resembling those triggered by IES (31,51,64).

### Aligning IES and TMS

The OFF periods triggered by IES during wakefulness did not seem to obliterate the emergence of complex interactions beyond the immediate area of stimulation—as evidenced by the higher PCI^ST^ values recorded during wakefulness compared to NREM sleep. The present findings help reconcile the apparently conflicting views on IES. While some studies used IES as a tool to probe global network connectivity (8,12,65), others suggest that electrical stimulation disrupts cortico-cortical signal propagation by silencing the output of target neocortical area (66,67). Our findings suggest that IEPs during wakefulness are characterized by a mixed response pattern wherein a strong local inhibition does not necessarily prevent the build-up of network interactions beyond the stimulated site. This is consistent with previous stereo-EEG evidence that IES during wakefulness triggers a stereotypical slow wave and an OFF-period near the stimulated site, while remote contacts show more complex responses (18). This pattern can be understood in the context of Fig. 5, assuming that areas that are distant from the site of IES receive a weaker activation that, similarly to TMS, does not necessarily engage activity-dependent mechanisms. Accordingly, the present finding that the complexity of IEPs exhibited state-dependent modulation between wakefulness and NREM sleep consistent with that observed in the case of TEPs, suggests that both stimulation techniques can induce large-scale responses that are informative about the global state of thalamocortical networks (35,68).

The differences and similarities between the effects of IES and TMS are also relevant to align and interpret responses to direct cortical stimulation across species and experimental models. IEPs have been recently used to replicate TMS-EEG results during wakefulness, sleep and anesthesia in rats (69,70), mice (70–72) and cortical slices (73). These works have shown consistent changes across global states but also differences in the profile of local activation, wherein slow responses and OFF-periods are often present also during wakefulness. In this context, the possibility that IES may saturate adaptation and inhibition mechanisms warrants caution and limits the interpretation of the literature, as area- or disease-specific reactivity profiles typically found in TEPs (74–76) may be masked or obliterated. It would be interesting to explore the parameter space of intracranial stimulation to attenuate the recruitment of activity-dependent mechanisms and match the local effects of TMS. This could potentially be obtained by lowering stimulation intensity while increasing the number of trials, and/or by diluting the E-field within a larger volume by stimulating across contacts farther apart instead of adjacent ones (77,78). Such exploration would also provide key empirical data to better understand the general activation mechanisms of TMS and their exquisite sensitivity in detecting changes in the state of cortical circuits (23,51,64,79).

### Limitations

Our primary goal was to understand the extent to which the effects of TMS and IES diverge under typical research and clinical conditions. Although we focused on relevant biophysical differences between the two techniques and on changes in brain states, it is important to recognize that other factors may have contributed to the present results.

Specifically, a potential limitation of this study is that TMS-EEG and IES-EEG data were collected from different populations: healthy participants and drug-resistant epileptic patients. In the latter, epilepsy might affect cortical networks near the epileptogenic zone. To mitigate this, we (i) included only patients without severe EEG abnormalities or malformations (80,81), (ii) avoided stimulating areas near the epileptogenic zone and excluded sessions with abnormal intracerebral responses (29), and (iii) confirmed that EEG responses to somatosensory stimulation were similar to those recorded in healthy subjects. Despite these precautions the authors acknowledge that differences in cortical health, disease burden, and medication status may contribute to the observed effects. Accordingly, results should be interpreted as reflecting the combined influences of stimulation modality and possibly population-level factors. Indeed the existing literature, although limited, demonstrates persistent differences in cortical reactivity to TMS between healthy subjects and epileptic patients (82,83). Yet, the differences observed between TMS and IES (e.g. evoked potentials ten times larger) cannot be fully explained by difference in cortical reactivity and are conceivably due mostly to stimulation modality. Future studies combining TMS-EEG and IES in the same patients (84) will further validate our findings.

Although TMS and IES targeted a common subset of cortical areas (e.g. Brodmann Areas 6, 7, 19), IES reached a broader set of more distributed regions, including deep structures like the insula and cingulate cortex. However, here we focused on general global (overall amplitude and complexity) or local (bistability) responsiveness brain features, which were previously shown to be independent on the specific site of stimulation (6,31,48,49). Additionally, we excluded areas causing motor or perceptual activations to avoid secondary EEG responses (48,85).

## Supplementary material

Supplementary material can be found in the online publication of this article.

## Author contribution

Conceptualization: MM, AP, EM, RC; Data curation: RC, AP, EM, SC, GH; Formal analysis: RC, GH, EM, MC; Funding acquisition: MM, AP; Investigation: SC, MF, GH, GF, EL, IS, AP, EM, SR, SP; Methodology: RC, EM, MC, SR, SD, EL; Project administration: MM, AP; Resources: MM; Software: RC, MC, SC, MF; Supervision: MM, AP; Validation; Visualization: RC, EM, AP, MC, SD; Writing – original draft: RC, MM, AP, SR; Writing – review and editing: MM, AP, RC, SR, EM, SC, MC.

## Acknowledgements

This work was supported by the European Research Council (ERC-2022-SYG - 101071900 - NEMESIS), and by the Ministry of University and Research (MUR), National Recovery and Resilience Plan (NRRP), project EBRAINS-Italy (IR00011). AP is supported by HORIZON-INFRA-2022 SERV (Grant No.101147319) “EBRAINS 2.0: A Research Infrastructure to Advance Neuroscience and Brain Health” and by Progetto Di Ricerca Di Rilevante Interesse Nazionale (PRIN) P2022FMK77. MM is supported by the ‘Brain, Mind, and Consciousness’ program of the Canadian Institute for Advanced Research (CIFAR). SD is supported by the Italian Ministry of Health – RicercaCorrente 2024.

