## Supplementart Materials for "Transcranial magnetic vs intracranial electric stimulation: a direct comparison of their effects via scalp EEG recordings"

### Supplementary Methods

*Participants and data acquisition*

TMS-EEG

TMS pulses were administered with a focal figure-of-eight coil (mean/outer winding diameter 50/70 mm) connected to a Mobile Stimulator Unit (eXimia TMS Stimulator, Nexstim Ltd.) (Fig. 1A). TMS targets were identified using a Navigated Brain Stimulation (NBS) system (Nexstim Ltd.) on a T1-weighted magnetic resonance of the subject's brain. The left and right Premotor (BA06), Parietal (BA07) and Occipital (BA19) cortices were targeted for the wakefulness dataset, and Parietal (BA07) or Premotor (BA06) for the paired NREM sleep dataset. TMS-EEG recordings were obtained from TMS-compatible EEG amplifiers (64-channels, Ag-Cl electrodes); specifically, a Brain Products GmbH amplifier (with a sampling rate of 5000 Hz; hardware filtered at 1000 Hz) and a Nexstim amplifier (with a sampling rate of 1450 Hz; hardware filtered at 350 Hz). The acquisition reference and ground were placed on the forehead (over the frontal sinus, near Fpz), and input-impedance was kept below 10 kOhms. Participants were seated in a comfortable chair both during wakefulness and NREM sleep, wearing headphones playing a customized noise to mask the TMS discharge sound (1).

Stimulation intensity, coil orientation, and position were determined for each participant to maximize TMS impact on the cortex (minimum early peak-to-peak amplitude of 8 μV) and minimize twitches of scalp muscles, using the real-time TEP visualization tool described in (2).

IES-EEG

Bipolar single-pulse IES was administered through stereotactically implanted platinum-iridium multi–contact intracerebral electrodes (0.8 mm diameter; 2 mm contact length; 1.5 mm inter-contact distance; Dixi Medical) at locations outside the epileptic area (Fig. 1B). Recordings were simultaneously conducted using 256 channels high-density scalp EEG (Geodesic Sensor Net; HydroCel CleanLeads) sampled at 1000 Hz using an EGI NA-400 amplifier (Electrical Geodesics, Inc; Oregon, USA).

IES duration and intensity was determined in the clinical setting to be generally effective for inducing subcortical and cortico-cortical evoked potentials (CCEPs), while also minimizing electric after-discharges and seizures (3–5). Further details of acquisitions are described in (6).

*Data Preprocessing*

TMS-EEG

First, the stimulation artifact was removed by replacing the interval around the stimulation pulse (-2 to 5 ms) with a mirrored version of the baseline signal, followed by a moving average filter with a 4 ms time span. This was done only to BrainAmp recordings, as the Nexstim system employs a built-in "sample and hold" mechanism that automatically pauses recording 2 ms before the pulse and resumes 2 ms after, eliminating the need for additional artifact correction. Beyond this system-specific correction, all remaining preprocessing steps were identical for data recorded with both EEG systems.

Next, the hd-EEG data was high-pass filtered at 0.5 Hz (zero-phase shift Butterworth, 3rd order) and splitted into epochs of ±800 ms centered around the TMS pulse, and bad trials and channels were rejected by visual inspection. The resulting epochs were re-referenced to the average reference and baseline corrected. Independent Component Analysis (ICA) was performed (infomax using *runica* EEGLAB routine (9)) to remove electro-oculographic (EOG) and electromyographic (EMG) activity (the latter being prominent in TMS but not in IES; average of 13.8 components removed). To enhance the separation between artifactual and neurophysiological components, ICA was performed on a narrower high-pass filtered (3rd order Butterworth, 1 Hz) version of the data, the unmixing weights were applied to the original data, and the retained components were back-projected to the scalp (7). After ICA, the signal was low-pass filtered at 45 Hz (zero-phase shift Butterworth, 3rd order) and down-sampled to 1000 Hz. Finally, bad channels were interpolated using spherical splines, and epochs were cropped from -600 to 600 ms.

IES-EEG

First of all, for each subject we excluded from the analysis all the sessions that showed marked epileptic activity based exclusively on the data recorded at the intracranial EEG level and we used the same criteria as in (8). Specifically, stimulation sessions were analyzed if they: (i) targeted contacts outside the epileptogenic zone (as confirmed by post-surgical assessment), (ii) showed no spontaneous interictal activity, (iii) evoked no epileptic responses during wakefulness or NREM (9), (iv) elicited no muscle twitches, sensations, or cognitive/motor effects during presurgical evaluation, including single-pulse and 50 Hz repetitive stimulation (10) occurred during N3 sleep, and (vi) did not disrupt sleep depth, as confirmed by comparing pre- and post-stimulation scalp EEG power spectra.

IES-EEG data were processed using a pipeline analogous to that employed for TMS-EEG data as detailed in (3). First, channels and trials contaminated by noise, muscle activity or spontaneous interictal epileptic discharges were rejected using a semi-automatic procedure, manually verified by an expert electrophysiologist. Next, the stimulation artifact was removed using a Tukey filter (8), data were band-pass filtered (0.5-45 Hz, zero-phase shift Butterworth, 3rd order) and epoched from -300 to 700 ms around IES triggers. Rejected channels were interpolated using spherical splines. Finally, trials were re-referenced to the average across all contacts, baseline corrected, and ICA was applied to remove EOG components (same algorithm as TMS; average of 1.3 components removed). Finally, data were downsampled to 1000 Hz.

*Data Analysis*

$PCI^{ST}$

The Perturbational Complexity Index state-transition ($PCI^{ST}$) is a measure developed to quantify the spatiotemporal complexity of brain responses to direct cortical perturbations (11), and provides an alternative method to its original method, the Lempel-Ziv-based Perturbational Complexity Index (PCI) (12), enabling faster and broader applicability across different stimulation and recording modalities (11).

$PCI^{ST}$ is computed in two main steps, each capturing a different aspect of complexity. First, Singular Value Decomposition (SVD) is applied to the trial-averaged evoked response, extracting the principal components that account for at least 99% of the total response power. This step captures the spatial aspect of complexity, since the greater the number of distinct spatial components, the higher the spatial differentiation of the response.

The second step quantifies the temporal aspect of complexity, which involves applying recurrence quantification analysis (RQA) to each spatial component (13). For each retained principal component, a distance matrix is computed by evaluating the voltage differences between all time points. A thresholding procedure is then applied to these distance matrices to identify significant deviations from baseline activity, which correspond to state transitions (ST). The total number of significant state transitions (ΔNST) over the optimal threshold is quantified for each component. Finally, $PCI^{ST}$ is obtained by summing the maximized ΔNST across all principal components, reflecting the degree to which the perturbation evokes a spatially structured yet diverse temporal pattern. $PCI^{ST}$ can also be defined as the product between a spatial term (the number of selected principal components) and a temporal term (the average number of significant state transitions across components), such that it is high when a perturbation evoked responses that are both spatially diverse and temporally rich, and low otherwise.

*Statistical analysis*

To further validate our findings, we performed Linear Mixed Effects Model (LMM) analyses in R using the lmerTest package. These models account for differences in the number of stimulated sites per subject, which vary across conditions. For the first part of the analysis (TMS vs. IES; Fig. 1-2), LMMs were used with stimulation modality as a fixed effect and random intercepts per subject to account for repeated measures within individuals. In the second part (TMS vs. IES, Wake vs. NREM sleep; Fig. 4), we included an interaction term between stimulation modality and state to examine their combined effects while still controlling for subject-level variability. Post-hoc comparisons were conducted using Estimated Marginal Means (emmeans), with multiple comparisons adjusted using Tukey's method where applicable. Model appropriateness was assessed via visual inspection of simulated residuals using the DHARMa package, ensuring assumptions were met. Results are reported in Tables S3-13.

#### Supplementary Figures

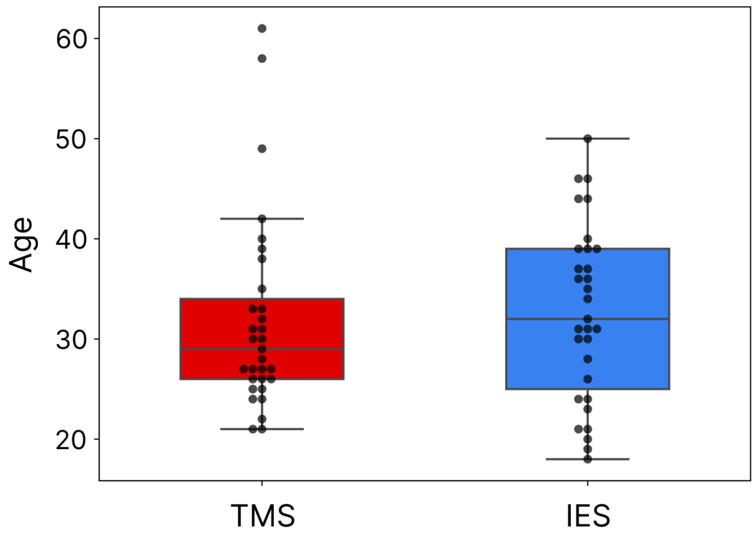


**Figure S1. Age distribution comparison between TMS and IES groups.** Boxplots display the age distribution for all participants in the TMS (blue) and IES (red) datasets. A Mann-Whitney U test revealed no significant difference between the two groups (U = 427.0000, p = 0.4551), confirming that they are age-matched.

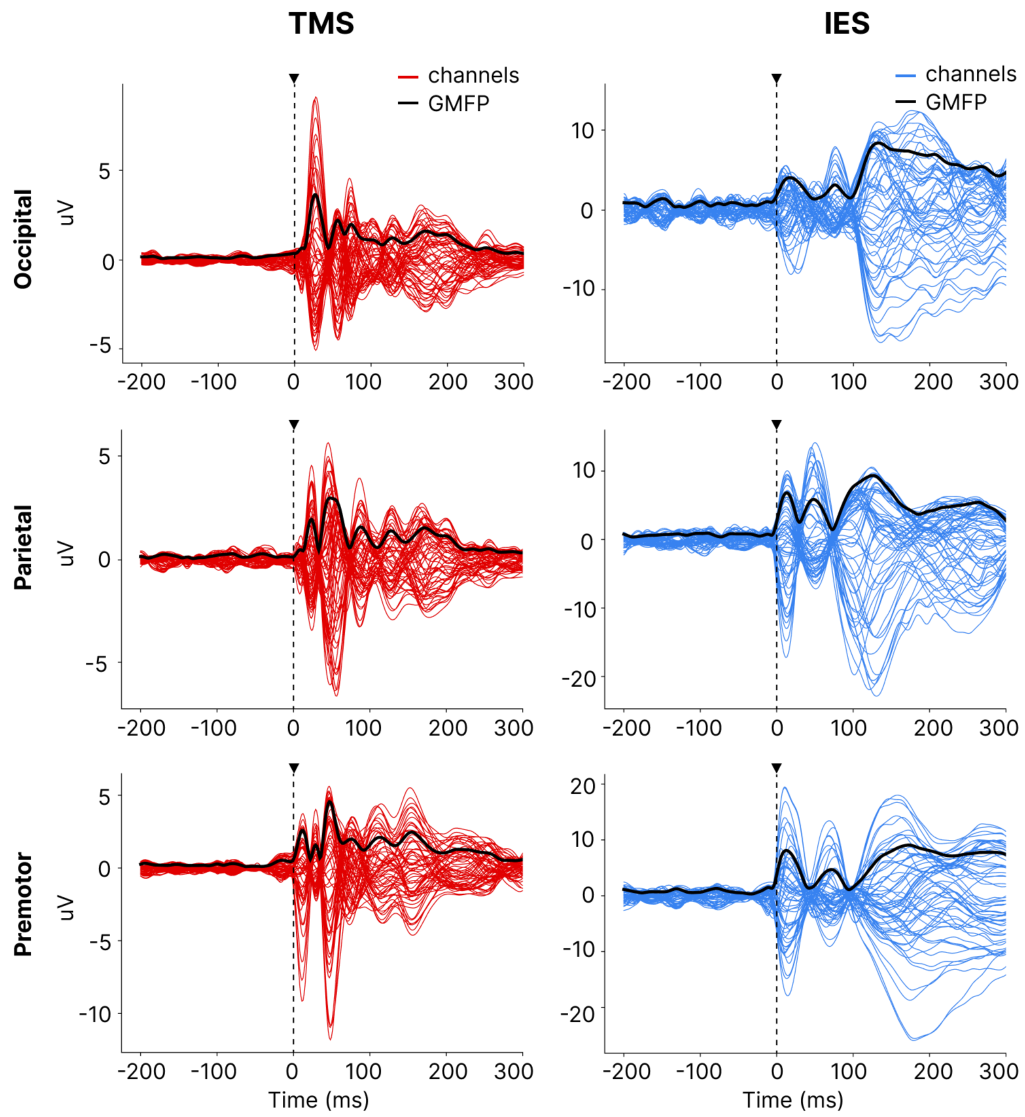


**Fig. S2. Representative TMS and IES evoked potentials.** (A) Butterfly plots of individual TEPs and IEPs obtained by the stimulation of occipital, parietal, and premotor areas in representative subjects. The colored traces depict the evoked response across EEG channels, with the GMFP overlaid in black, illustrating the power evoked globally across the EEG channels over time.

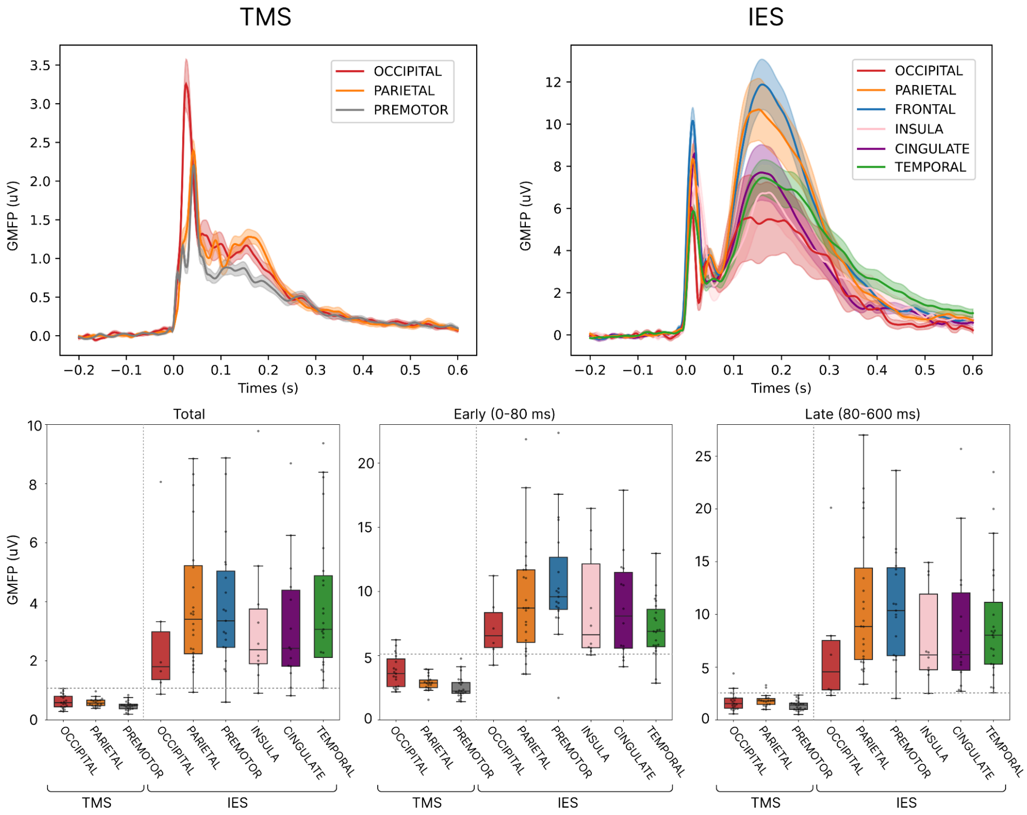


**Fig. S3. Site-specificity of TMS and IES evoked responses.** (A) Global Mean Field Power (GMFP) time courses of EEG responses evoked by TMS (left) and IES (right) for different stimulation sites. Colored traces represent the GMFP for each cortical region, with shaded areas indicating the standard error of the mean. TMS responses (left) are shown for stimulation of the occipital (red), parietal (orange), and premotor (gray) cortices. IES responses (right) are shown for occipital (red), parietal (blue), frontal (orange), insula (purple), cingulate (pink), and temporal (green) cortices. (B) Boxplots of GMFP amplitudes comparing TMS and IES responses across stimulation sites. Three response windows are considered: total response (0–600 ms, left panel), early response (0–80 ms, middle panel), and late response (80–600 ms, right panel). Each boxplot shows the median, interquartile range, and individual data points.

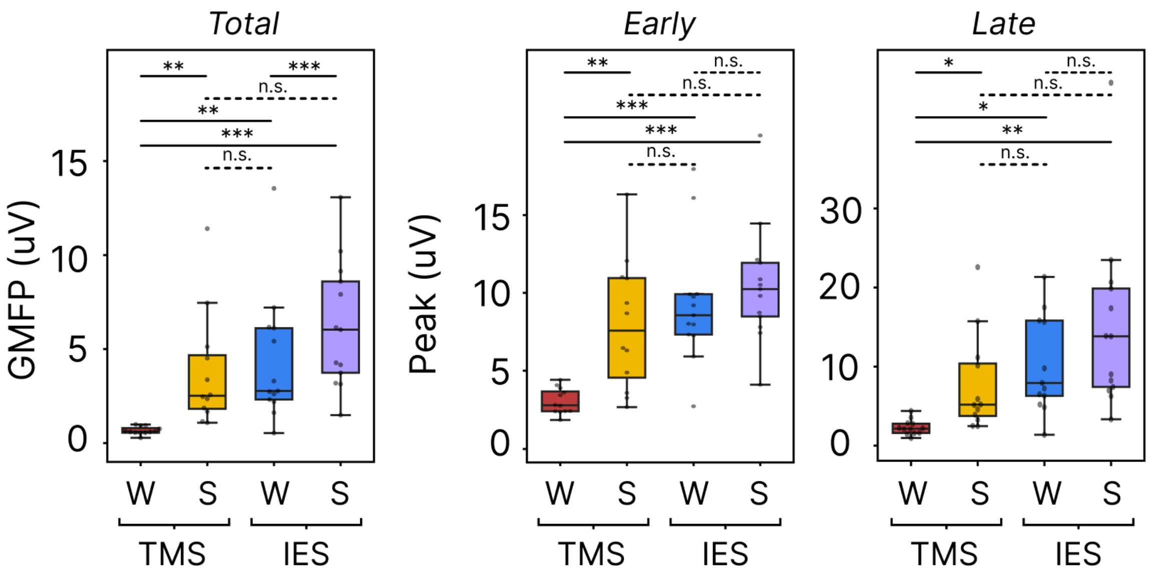


**Fig. S4. Global Mean Field Power (GMFP) metrics across wakefulness and NREM sleep for TMS and IES.** Boxplots showing GMFP across the full response interval (0-600 ms), and the peak GMFP amplitudes for the early (0-80 ms) and late (80-600 ms) intervals, for TMS and IES during wakefulness (W) and NREM sleep (S). Each boxplot displays the median, interquartile range, and individual data points. Statistical significance is denoted by asterisks (n.s. = $p > 0.05$, * = $p < 0.05$, ** = $p < 0.01$, *** = $p < 0.001$).

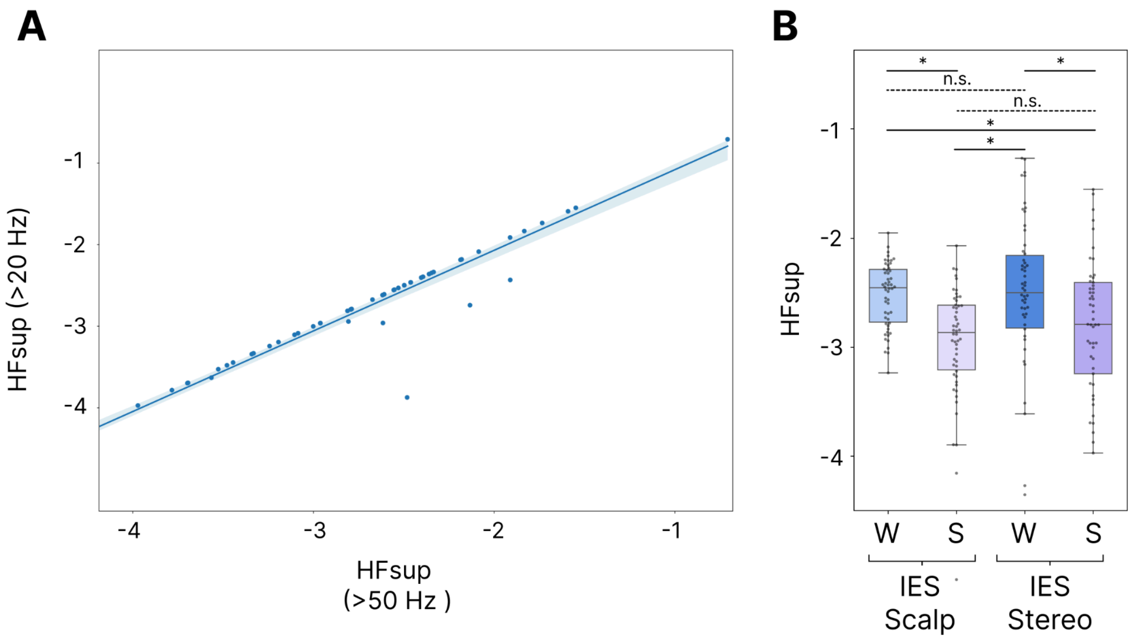


**Fig. S5. Comparison of high-frequency suppression across different frequency ranges (>20 Hz vs. >50 Hz) and spatial scales (EEG vs. SEEG).** Panel (A) shows, for all IEPs (each bullet), a comparison of High-Frequency Suppression (HFsup) calculated from two different frequency ranges (>20 Hz and >50 Hz) using the SEEG contact located directly beneath the scalp EEG electrode that exhibited the largest response to IES. Panel (B) compares HFsup (>20 Hz) between the scalp EEG electrode showing the strongest response to IES and the SEEG contact positioned directly beneath it (i.e., the closest in geometric proximity), for each IEP (n.s. = $p > 0.05$, * = $p < 0.05$, ** = $p < 0.01$, *** = $p < 0.001$).

#### Supplementary Tables

| Subject | Sex | Age | States | Stimulated areas (hemisphere) |
|---|---|---|---|---|
| **1** | F | 27 | W | occipital (L), parietal (L), premotor (L, R) |
| **2** | F | 22 | W | parietal (L), premotor (L) |
| **3** | M | 32 | W | occipital (L, R), premotor (L, R) |
| **4** | F | 61 | W | occipital (L, R), parietal (L, R), premotor (L, R) |
| **5** | M | 27 | W | occipital (L), parietal (L), premotor (L, R) |
| **6** | M | 40 | W | occipital (R) |
| **7** | M | 26 | W | occipital (L, R), parietal (L, R), premotor (L, R) |
| **8** | M | 25 | W | occipital (L, R), premotor (L) |
| **9** | M | 28 | W | occipital (L), parietal (L), premotor (L) |
| **10** | F | 27 | W | occipital (L, R), parietal (L, R), premotor (L, R) |
| **11** | F | 27 | W | occipital (L), parietal (L, R), premotor (L, R) |
| **12** | F | 24 | W | occipital (L, R), parietal (L), premotor (L) |
| **13** | F | 31 | W | occipital (L), parietal (L), premotor (L) |
| **14** | F | 25 | W | occipital (L), parietal (L), premotor (L, R) |
| **15** | F | 24 | W | occipital (L), parietal (L), premotor (L) |
| **16** | F | 30 | W | premotor (L) |
| **17** | M | 33 | W | occipital (L, R), parietal (L), premotor (L, R) |
| **18** | F | 58 | W | occipital (L, R), parietal (L, R), premotor (L, R) |
| **19** | M | 26 | W | occipital (L), parietal (L), premotor (L, R) |
| **20** | F | 26 | W, S | occipital (L, R), parietal (L*, R), premotor (L, R) |
| **21** | F | 29 | W, S | parietal (L*, R), premotor (L, R) |

**Table S1: Demographic and stimulation information for each TMS/EEG subject.**
F: female; M: male; W: wakefulness; S: NREM sleep; L: left, R: right

| Patient | Sex | Age | MRI | Hemisphere of SEEG | Lobes of SEEG | Therapy |
|---|---|---|---|---|---|---|
| **1** | F | 31 | Negative | L | temporo-parieto-perisy lvian | BRV 200 mg/die; PER 6 mg/die |
| **2** | M | 21 | Negative | L | frontal | CBZ 1400 mg/die; PB 50 mg/die; CLB 10 mg/die |
| **3** | F | 26 | Negative | R | fronto-centro-parieto-t emporo-perisylvian | LEV 3000 mg/die; TPM 300 mg/die |
| **4** | M | 39 | Negative | L | temporo-occipito-parie tal | CBZ 1600 mg/die; CLB 20 mg/die |
| **5** | M | 46 | Negative | L | temporo-parieto-perisy lvian | LTG 350 mg/die; OXC 1500 mg/die |
| **6** | F | 30 | Negative | L | fronto-temporal | CBZ 1000 mg/die; LTG 625 mg/die; CLB 20 mg/die; PGB 20 mg/die |
| **7** | M | 18 | Negative | L | fronto-central | LCM 500 mg/die; TPM 200 mg/die |
| **8** | M | 44 | Negative | Bilat | Right temporo-occipito-parietal + 2 electrodes in left temporal lobe | CBZ 1200 mg/die, PMP 6 mg/die |
| **9** | F | 21 | Negative | L | perisylvian | CBZ 500 mg/die; CLB 20 mg/die; ESL 1200 mg/die; PER 8 mg/die |
| **10** | F | 39 | Negative | L | temporo-parieto-perisy lvian | CBZ 1000 mg/die; LEV 2750 mg/die; CLB 10 mg/die |
| **11** | M | 19 | Negative | R | temporo-occipito-parie tal | CBZ 1800 mg/die; LCM 200 mg /die |
| **12** | F | 50 | Negative | L | temporo-occipito-parie tal | LCM 400 mg/die; ZNS 400 mg/die; PB 100 mg/die |
| **13** | F | 20 | Negative | L | temporo-parieto-perisy lvian | LTG 200 mg/die; CLB 10 mg/die |
| **14** | F | 28 | Negative | Bilat | Right temporo-perisylvian+left temporo-fronto-perisylvian | LCM 400 mg/die; OXC 600 mg/die |
| **15** | F | 24 | Negative | R | fronto-centro-parietal | VPA 1500 mg/die; LCM 400 mg/die |
| **16** | M | 37 | Negative | R | fronto-centro-parieto-perysilvian | CBZ 800 mg/die; LCM 400 mg/die |
| **17** | M | 36 | Negative | L | temporo- perisylvian | LEV 3000; LCM 400 mg/die; CBZ 1400 mg/die |
| **18** | M | 37 | Negative | R | fronto-central + 1 electrode in parietal lobe + | CBZ 1600 mg/die; TPM 300 mg/die |

| | | | | | | |
|---|---|---|---|---|---|---|
| | | | | | 1 electrode in temporal lobe (hipp) | |
| **19** | F | 40 | Negative | L | temporo-occipito-parietal | LTG 400 mg/die; TPM 100 mg/die |
| **20** | M | 35 | Negative | R | fronto-centro-temporal | CBZ 1400 mg/die; LEV 750 mg/die |
| **21** | F | 31 | Negative | Bilat | bilateral temporo-perisylvian (> on the right) | CBZ 1200 mg/die; TPM 150 mg/die |
| **22** | F | 32 | Negative | Bilat | Right temporo-occipito-parietal-perysilvian+3 electrodes on left temporal lobe | CBZ 800 mg/die; ZNS 200 mg/die; CLB 20 mg/die |
| **23** | F | 34 | Negative | L | temporo-perisylvian | CBZ 1600 mg/die; PB 100 mg/die |
| **24** | M | 30 | Negative | Bilat | Fronto-central (> on the left) | OXC 1800 mg/die; TPM 200 mg/die; LEV 3000 mg/die; CLB 10 mg/die. |
| **25** | F | 39 | Negative | Bilat | Right temporo-perysislvio-central + left temporo-perysislvian (> a destra) | LTG 400 mg/die; PMP 8 mg/die |
| **26** | F | 31 | Negative | L | temporo-occipito-parietal | LTG 6000 mg/die; ZNS 450 mg/die; CLB 50 mg/die; CLZ 3 mg |
| **27** | M | 23 | Negative | L | temporo-perisylvian | ZNS 200 mg/die; PB 150 mg/die |
| **28** | M | 44 | Negative | R | temporo-occipito-parietal | CBZ 1400 mg/die; PB 100 mg/die; LTG 400 mg/die |
| **29** | F | 24 | Negative | R | temporo-occipito-parietal | CBZ 1500 mg/die; TPM 75 mg/die |
| **30** | F | 46 | Negative | L | temporo-perisylvian | OXC 1200 mg/die; LCS 150 mg/die |
| **31** | F | 36 | Negative | L | front-temporo-perisylvian | LTG 400 mg/die; CBZ 1200 mg/die |

**Table S2: Demographic and clinical details for each patient**

BRV: Brivaracetam; CBZ: Carmabazepine; CLB: Clobazam; CLZ: Clonazepam; ESL: Eslicarbazepine acetate; LCM: Lacosamide; LEV: Levetiracetam; LTG: Lamotrigine; OXC: Oxcarbazepine; PB: Phenobarbital; PER: Perampanel; PGB: Pregabalin; TPM: Topiramate; VPA: Valproate; ZNS: Zonisamide

```
Linear mixed model fit by REML. t-tests use Satterthwaite's method ['lmerModLmerTest']
Formula: ntrials ~ stimulation + (1 | subj)
   Data: dat_all

REML criterion at convergence: 2845.4

Scaled residuals: 
    Min      1Q  Median      3Q     Max 
-5.0344 -0.0702 -0.0130  0.0991  8.2303 

Random effects:
 Groups   Name        Variance Std.Dev.
 subj     (Intercept) 142.1    11.92   
 Residual             352.4    18.77   
Number of obs: 321, groups:  subj, 53

Fixed effects:
               Estimate Std. Error      df t value Pr(>|t|)    
(Intercept)      32.040      2.538  28.395   12.62 3.62e-13 ***
stimulationtms  196.132      4.156  35.284   47.20  < 2e-16 ***
---
Signif. codes:  0 '***' 0.001 '**' 0.01 '*' 0.05 '.' 0.1 ' ' 1

Correlation of Fixed Effects:
            (Intr)
stimultntms -0.611
```

#### Table S3: Linear Mixed Model for number of trials (Fig 1E)

```
Linear mixed model fit by REML. t-tests use Satterthwaite's method ['lmerModLmerTest']
Formula: snr_20_trials ~ stimulation + (1 | subj)
   Data: dat_all

REML criterion at convergence: 1973

Scaled residuals: 
    Min      1Q  Median      3Q     Max 
-1.9547 -0.5663 -0.1960  0.3825  5.3399 

Random effects:
 Groups   Name        Variance Std.Dev.
 subj     (Intercept)  7.61    2.759   
 Residual             24.94    4.994   
Number of obs: 318, groups:  subj, 53

Fixed effects:
               Estimate Std. Error      df t value Pr(>|t|)    
(Intercept)     11.2543     0.6134 48.0293  18.347  < 2e-16 ***
stimulationtms  -5.8299     1.0131 60.7525  -5.755 3.04e-07 ***
---
Signif. codes:  0 '***' 0.001 '**' 0.01 '*' 0.05 '.' 0.1 ' ' 1

Correlation of Fixed Effects:
            (Intr)
stimultntms -0.606
```

#### Table S4: Linear Mixed Model for SNR (20 trials) (Fig 1E)

```
Linear mixed model fit by REML. t-tests use Satterthwaite's method ['lmerModLmerTest']
Formula: snr_all_trials ~ stimulation + (1 | subj)
   Data: dat_all

REML criterion at convergence: 2230.3

Scaled residuals: 
    Min      1Q  Median      3Q     Max 
-1.7595 -0.6648 -0.2166  0.4267  4.9039 

Random effects:
```

```
 Groups   Name        Variance Std.Dev.
 subj     (Intercept) 14.61    3.822
 Residual             53.24    7.297
Number of obs: 321, groups:  subj, 53

Fixed effects:
               Estimate Std. Error      df t value Pr(>|t|)
(Intercept)     13.6910     0.8623 39.5604  15.877   <2e-16 ***
stimulationtms   2.3679     1.4329 51.6745   1.653    0.104
---
Signif. codes:  0 '***' 0.001 '**' 0.01 '*' 0.05 '.' 0.1 ' ' 1

Correlation of Fixed Effects:
            (Intr)
stimultntms -0.602
```

##### Table S5: Linear Mixed Model for SNR (all trials) (Fig 1E)

```
Linear mixed model fit by REML. t-tests use Satterthwaite's method ['lmerModLmerTest']
Formula: gmfp_total ~ stimulation + (1 | subj)
   Data: dat_all

REML criterion at convergence: -7328.1

Scaled residuals:
    Min      1Q  Median      3Q     Max
-2.4913 -0.4617 -0.0427  0.1310  4.2877

Random effects:
 Groups   Name        Variance  Std.Dev.
 subj     (Intercept) 2.056e-12 1.434e-06
 Residual             4.958e-12 2.227e-06
Number of obs: 321, groups:  subj, 53

Fixed effects:
                 Estimate Std. Error         df t value Pr(>|t|)
(Intercept)     3.797e-06  3.041e-07  1.202e+01  12.486 3.05e-08 ***
stimulationtms -3.240e-06  4.975e-07  1.255e+01  -6.512 2.33e-05 ***
---
Signif. codes:  0 '***' 0.001 '**' 0.01 '*' 0.05 '.' 0.1 ' ' 1

Correlation of Fixed Effects:
            (Intr)
stimultntms -0.611
```

##### Table S6: Linear Mixed Model for GMFP Total (Fig 2C)

```
Linear mixed model fit by REML. t-tests use Satterthwaite's method ['lmerModLmerTest']
Formula: gmfp_early_peak ~ stimulation + (1 | subj)
   Data: dat_all

REML criterion at convergence: -7027.2

Scaled residuals:
    Min      1Q  Median      3Q     Max
-2.9296 -0.5784 -0.0964  0.3784  4.5504

Random effects:
 Groups   Name        Variance  Std.Dev.
 subj     (Intercept) 3.898e-12 1.974e-06
 Residual             1.318e-11 3.630e-06
Number of obs: 321, groups:  subj, 53

Fixed effects:
                 Estimate Std. Error         df t value Pr(>|t|)
(Intercept)     8.719e-06  4.399e-07  1.687e+01   19.82 4.01e-13 ***
stimulationtms -5.649e-06  7.288e-07  1.816e+01   -7.75 3.62e-07 ***
---
Signif. codes:  0 '***' 0.001 '**' 0.01 '*' 0.05 '.' 0.1 ' ' 1
```

```
Correlation of Fixed Effects:
            (Intr)
stimultntms -0.604
```

#### Table S7: Linear Mixed Model for GMFP Early Peak (Fig 2C)

```
Linear mixed model fit by REML. t-tests use Satterthwaite's method ['lmerModLmerTest']
Formula: gmfp_late_peak ~ stimulation + (1 | subj)
   Data: dat_all

REML criterion at convergence: -6669.7

Scaled residuals: 
    Min      1Q  Median      3Q     Max 
-2.4659 -0.4454 -0.0659  0.1645  5.9063 

Random effects:
 Groups   Name        Variance  Std.Dev. 
 subj     (Intercept) 1.677e-11 4.095e-06
 Residual             3.890e-11 6.237e-06
Number of obs: 321, groups:  subj, 53

Fixed effects:
                 Estimate Std. Error         df t value Pr(>|t|)    
(Intercept)     1.016e-05  8.643e-07  2.176e+01  11.758 6.76e-11 ***
stimulationtms -8.535e-06  1.412e-06  2.311e+01  -6.046 3.56e-06 ***
---
Signif. codes:  0 '***' 0.001 '**' 0.01 '*' 0.05 '.' 0.1 ' ' 1

Correlation of Fixed Effects:
            (Intr)
stimultntms -0.612
```

#### Table S8: Linear Mixed Model for GMFP Late Peak (Fig 2C)

*With interaction*

```
Linear mixed model fit by REML. t-tests use Satterthwaite's method ['lmerModLmerTest']
Formula: gmfp_total ~ stimulation * state + (1 | subj)
   Data: dat

REML criterion at convergence: 643.4

Scaled residuals: 
    Min      1Q  Median      3Q     Max 
-2.4185 -0.6235 -0.1015  0.4350  3.1384 

Random effects:
 Groups   Name        Variance Std.Dev.
 subj     (Intercept) 4.724    2.174   
 Residual             6.281    2.506   
Number of obs: 132, groups:  subj, 25

Fixed effects:
                           Estimate Std. Error       df t value Pr(>|t|)   
(Intercept)                  0.6671     0.9577  58.3323   0.697  0.48880   
stimulationseeg              3.5574     1.1932  39.9411   2.981  0.00487 **
statesleep                   3.0836     1.0232 105.5257   3.014  0.00323 **
stimulationseeg:statesleep  -0.9182     1.1311 105.5257  -0.812  0.41879   
---
Signif. codes:  0 '***' 0.001 '**' 0.01 '*' 0.05 '.' 0.1 ' ' 1
Correlation of Fixed Effects:
            (Intr) stmltn sttslp
stimulatnsg -0.803              
statesleep  -0.534  0.429       
stmltnsg:st  0.483 -0.474 -0.905

$emmeans
 stimulation state emmean    SE   df lower.CL upper.CL
 tms         wake   0.667 0.958 58.8    -1.25     2.58
 seeg        wake   4.224 0.713 22.3     2.75     5.70
```

```
 tms         sleep  3.751 0.958 58.8     1.83     5.67
 seeg        sleep  6.390 0.713 22.3     4.91     7.87

Degrees-of-freedom method: kenward-roger
Confidence level used: 0.95

$contrasts
 contrast                estimate    SE    df t.ratio p.value
 tms wake - seeg wake      -3.557 1.194  40.4  -2.979  0.0242
 tms wake - tms sleep      -3.084 1.023 105.8  -3.014  0.0168
 tms wake - seeg sleep     -5.723 1.194  40.4  -4.792  0.0001
 seeg wake - tms sleep      0.474 1.194  40.4   0.397  0.9785
 seeg wake - seeg sleep    -2.165 0.482 105.8  -4.490  0.0001
 tms sleep - seeg sleep    -2.639 1.194  40.4  -2.210  0.1378

Degrees-of-freedom method: kenward-roger
P value adjustment: tukey method for comparing a family of 4 estimates
```

*Without interaction*

```
Linear mixed model fit by REML. t-tests use Satterthwaite's method ['lmerModLmerTest']
Formula: gmfp_total ~ stimulation + state + (1 | subj)
   Data: dat

REML criterion at convergence: 646.1

Scaled residuals:
    Min      1Q  Median      3Q     Max
-2.4560 -0.5982 -0.1441  0.4531  3.1764

Random effects:
 Groups   Name        Variance Std.Dev.
 subj     (Intercept) 4.730    2.175
 Residual             6.261    2.502
Number of obs: 132, groups:  subj, 25

Fixed effects:
                Estimate Std. Error       df t value Pr(>|t|)
(Intercept)       1.0427     0.8381  37.0051   1.244  0.22128
stimulationseeg   3.0983     1.0506  24.6094   2.949  0.00689 **
statesleep        2.3324     0.4356 106.5154   5.355 4.97e-07 ***
---
Signif. codes:  0 '***' 0.001 '**' 0.01 '*' 0.05 '.' 0.1 ' ' 1

Correlation of Fixed Effects:
            (Intr) stmltn
stimulatnsg -0.744
statesleep  -0.260  0.000

$emmeans
 stimulation state emmean    SE   df lower.CL upper.CL
 tms         wake    1.04 0.838 37.4   -0.655     2.74
 seeg        wake    4.14 0.706 21.4    2.675     5.61
 tms         sleep   3.38 0.838 37.4    1.678     5.07
 seeg        sleep   6.47 0.706 21.4    5.007     7.94

Degrees-of-freedom method: kenward-roger
Confidence level used: 0.95

$contrasts
 contrast                estimate    SE    df t.ratio p.value
 tms wake - seeg wake      -3.098 1.052  24.9  -2.946  0.0326 *
 tms wake - tms sleep      -2.332 0.436 106.8  -5.355  <.0001 ***
 tms wake - seeg sleep     -5.431 1.138  33.8  -4.771  0.0002 ***
 seeg wake - tms sleep      0.766 1.138  33.8   0.673  0.9066
 seeg wake - seeg sleep    -2.332 0.436 106.8  -5.355  <.0001 ***
 tms sleep - seeg sleep    -3.098 1.052  24.9  -2.946  0.0326 *

Degrees-of-freedom method: kenward-roger
P value adjustment: tukey method for comparing a family of 4 estimates
```

**Table S9: Linear Mixed Model for GMFP (wake/sleep) (Fig 4C)**

```
Linear mixed model fit by REML. t-tests use Satterthwaite's method ['lmerModLmerTest']
Formula: pci ~ stimulation * state + (1 | subj)
   Data: dat

REML criterion at convergence: 989.9

Scaled residuals: 
    Min      1Q  Median      3Q     Max 
-2.3864 -0.5848 -0.0663  0.5456  2.4790 

Random effects:
 Groups   Name        Variance Std.Dev.
 subj     (Intercept)  42.73    6.537  
 Residual             100.65   10.033  
Number of obs: 132, groups:  subj, 25

Fixed effects:
                           Estimate Std. Error      df t value Pr(>|t|)    
(Intercept)                  65.871      3.457  82.087  19.056  < 2e-16 ***
stimulationseeg             -28.876      4.177  55.099  -6.913 5.20e-09 ***
statesleep                  -41.351      4.096 109.051 -10.096  < 2e-16 ***
stimulationseeg:statesleep   26.383      4.528 109.051   5.827 5.82e-08 ***
---
Signif. codes:  0 '***' 0.001 '**' 0.01 '*' 0.05 '.' 0.1 ' ' 1

Correlation of Fixed Effects:
            (Intr) stmltn sttslp
stimulatnsg -0.828              
statesleep  -0.592  0.490       
stmltnsg:st  0.536 -0.542 -0.905

$emmeans
 stimulation state emmean   SE   df lower.CL upper.CL
 tms         wake    65.9 3.46 77.9     59.0     72.8
 seeg        wake    37.0 2.35 22.4     32.1     41.9
 tms         sleep   24.5 3.46 77.9     17.6     31.4
 seeg        sleep   22.0 2.35 22.4     17.1     26.9

Degrees-of-freedom method: kenward-roger 
Confidence level used: 0.95 

$contrasts
 contrast               estimate   SE    df t.ratio p.value
 tms wake - seeg wake      28.88 4.18  50.8   6.904  <.0001 ***
 tms wake - tms sleep      41.35 4.10 106.7  10.096  <.0001 ***
 tms wake - seeg sleep     43.84 4.18  50.8  10.483  <.0001 ***
 seeg wake - tms sleep     12.47 4.18  50.8   2.982  0.0221 *
 seeg wake - seeg sleep    14.97 1.93 106.7   7.752  <.0001 ***
 tms sleep - seeg sleep     2.49 4.18  50.8   0.596  0.9328

Degrees-of-freedom method: kenward-roger 
P value adjustment: tukey method for comparing a family of 4 estimates
```

#### Table S10: Linear Mixed Model for PCIst (wake/sleep) (Fig 4C)

```
Linear mixed model fit by REML. t-tests use Satterthwaite's method ['lmerModLmerTest']
Formula: HFp_avgmin ~ stimulation * state + (1 | subj)
   Data: dat

REML criterion at convergence: 273.9

Scaled residuals: 
    Min      1Q  Median      3Q     Max 
-3.3553 -0.4887  0.0363  0.3475  6.4476 

Random effects:
 Groups   Name        Variance Std.Dev.
 subj     (Intercept) 0.000154 0.01241 
 Residual             0.449629 0.67054 
Number of obs: 132, groups:  subj, 25
```

```
Fixed effects:
                            Estimate Std. Error        df t value Pr(>|t|)
(Intercept)                 -0.09676    0.19360 127.94588  -0.500 0.618086
stimulationseeg             -2.36034    0.21406 120.41306 -11.026  < 2e-16 ***
statesleep                  -1.09130    0.27375 119.15840  -3.987 0.000116 ***
stimulationseeg:statesleep   0.66480    0.30264 119.15840   2.197 0.029980 *
---
Signif. codes:  0 '***' 0.001 '**' 0.01 '*' 0.05 '.' 0.1 ' ' 1

Correlation of Fixed Effects:
            (Intr) stmltn sttslp
stimulatnsg -0.904
statesleep  -0.707  0.639
stmltnsg:st  0.639 -0.707 -0.905

$emmeans
 stimulation state  emmean     SE    df lower.CL upper.CL
 tms         wake  -0.0968 0.1936 127.9    -0.48    0.286
 seeg        wake  -2.4571 0.0926  35.3    -2.64   -2.269
 tms         sleep -1.1881 0.1936 127.9    -1.57   -0.805
 seeg        sleep -2.8836 0.0926  35.3    -3.07   -2.696

Degrees-of-freedom method: kenward-roger
Confidence level used: 0.95

$contrasts
 contrast                estimate    SE  df t.ratio p.value
 tms wake - seeg wake       2.360 0.215 116  10.999  <.0001 ***
 tms wake - tms sleep       1.091 0.274 114   3.987  0.0007 ***
 tms wake - seeg sleep      2.787 0.215 116  12.987  <.0001 ***
 seeg wake - tms sleep     -1.269 0.215 116  -5.914  <.0001 ***
 seeg wake - seeg sleep     0.427 0.129 114   3.305  0.0069 **
 tms sleep - seeg sleep     1.696 0.215 116   7.901  <.0001 ***

Degrees-of-freedom method: kenward-roger
P value adjustment: tukey method for comparing a family of 4 estimates
```

##### Table S11: Linear Mixed Model for max HFsup (wake/sleep) (Fig 4F)

```
Linear mixed model fit by REML. t-tests use Satterthwaite's method ['lmerModLmerTest']
Formula: HFp_percent ~ stimulation * state + (1 | subj)
   Data: dat

REML criterion at convergence: -54.7

Scaled residuals:
    Min      1Q  Median      3Q     Max
-2.1357 -0.7113 -0.1041  0.5468  3.4976

Random effects:
 Groups   Name        Variance Std.Dev.
 subj     (Intercept) 0.004503 0.0671
 Residual             0.031566 0.1777
Number of obs: 132, groups:  subj, 25

Fixed effects:
                            Estimate Std. Error        df t value Pr(>|t|)
(Intercept)                  0.04727    0.05482 116.32166   0.862 0.390380
stimulationseeg              0.21592    0.06322  88.76615   3.415 0.000963 ***
statesleep                   0.41223    0.07253 114.38879   5.683 1.02e-07 ***
stimulationseeg:statesleep  -0.33026    0.08019 114.38879  -4.119 7.23e-05 ***
---
Signif. codes:  0 '***' 0.001 '**' 0.01 '*' 0.05 '.' 0.1 ' ' 1

Correlation of Fixed Effects:
            (Intr) stmltn sttslp
stimulatnsg -0.867
statesleep  -0.661  0.574
stmltnsg:st  0.598 -0.634 -0.905
```

```
$emmeans
 stimulation state emmean     SE    df lower.CL upper.CL
 tms         wake  0.0473 0.0548 112.1  -0.0614    0.156
 seeg        wake  0.2632 0.0317  24.7   0.1978    0.329
 tms         sleep 0.4595 0.0548 112.1   0.3509    0.568
 seeg        sleep 0.3452 0.0317  24.7   0.2797    0.411

Degrees-of-freedom method: kenward-roger
Confidence level used: 0.95

$contrasts
 contrast               estimate     SE    df t.ratio p.value
 tms wake - seeg wake     -0.216 0.0633  78.7  -3.408  0.0056 **
 tms wake - tms sleep     -0.412 0.0725 109.5  -5.683  <.0001 ***
 tms wake - seeg sleep    -0.298 0.0633  78.7  -4.702  0.0001 ***
 seeg wake - tms sleep    -0.196 0.0633  78.7  -3.099  0.0140 *
 seeg wake - seeg sleep   -0.082 0.0342 109.5  -2.397  0.0836
 tms sleep - seeg sleep    0.114 0.0633  78.7   1.805  0.2787

Degrees-of-freedom method: kenward-roger
P value adjustment: tukey method for comparing a family of 4 estimates
```

#### Table S12: Linear Mixed Model for %ch HFsup (wake/sleep) (Fig 4F)

*With Interaction*

```
Linear mixed model fit by REML. t-tests use Satterthwaite's method ['lmerModLmerTest']
Formula: HFp_negarea_int ~ stimulation * state + (1 | subj)
   Data: dat

REML criterion at convergence: 147.1

Scaled residuals: 
    Min      1Q  Median      3Q     Max 
-4.1671 -0.4039  0.0946  0.6793  2.0712 

Random effects:
 Groups   Name        Variance Std.Dev.
 subj     (Intercept) 0.01469  0.1212  
 Residual             0.15647  0.3956  
Number of obs: 132, groups:  subj, 25

Fixed effects:
                            Estimate Std. Error        df t value Pr(>|t|)
(Intercept)                 -0.02446    0.11943 122.77778  -0.205 0.838034
stimulationseeg             -0.49795    0.13607 102.17898  -3.659 0.000402 ***
statesleep                  -0.38113    0.16149 117.38035  -2.360 0.019920 *
stimulationseeg:statesleep   0.14372    0.17853 117.38035   0.805 0.422456
---
Signif. codes:  0 '***' 0.001 '**' 0.01 '*' 0.05 '.' 0.1 ' ' 1

Correlation of Fixed Effects:
            (Intr) stmltn sttslp
stimulatnsg -0.878              
statesleep  -0.676  0.593       
stmltnsg:st  0.612 -0.656 -0.905

$emmeans
 stimulation state  emmean     SE    df lower.CL upper.CL
 tms         wake  -0.0245 0.1194 119.2   -0.261    0.212
 seeg        wake  -0.5224 0.0658  26.2   -0.658   -0.387
 tms         sleep -0.4056 0.1194 119.2   -0.642   -0.169
 seeg        sleep -0.7598 0.0658  26.2   -0.895   -0.625

Degrees-of-freedom method: kenward-roger
Confidence level used: 0.95

$contrasts
 contrast               estimate     SE    df t.ratio p.value
 tms wake - seeg wake      0.498 0.1364  89.1   3.652  0.0024
```

```
 tms wake - tms sleep      0.381 0.1615 110.7   2.360  0.0910
 tms wake - seeg sleep     0.735 0.1364  89.1   5.393  <.0001
 seeg wake - tms sleep    -0.117 0.1364  89.1  -0.857  0.8269
 seeg wake - seeg sleep    0.237 0.0761 110.7   3.119  0.0122
 tms sleep - seeg sleep    0.354 0.1364  89.1   2.598  0.0526

Degrees-of-freedom method: kenward-roger
P value adjustment: tukey method for comparing a family of 4 estimates
```

*Without Interaction*

```
Linear mixed model fit by REML. t-tests use Satterthwaite's method ['lmerModLmerTest']
Formula: HFp_negarea_int ~ stimulation + state + (1 | subj)
   Data: dat

REML criterion at convergence: 146.2

Scaled residuals:
    Min      1Q  Median      3Q     Max
-4.1408 -0.4547  0.1645  0.6473  2.1076

Random effects:
 Groups   Name        Variance Std.Dev.
 subj     (Intercept) 0.01472  0.1213
 Residual             0.15601  0.3950
Number of obs: 132, groups:  subj, 25

Fixed effects:
                 Estimate Std. Error        df t value Pr(>|t|)
(Intercept)      -0.08326    0.09439 102.00395  -0.882 0.379799
stimulationseeg  -0.42607    0.10261  57.37981  -4.152 0.000110 ***
statesleep       -0.26354    0.06876 118.37151  -3.833 0.000204 ***
---
Signif. codes:  0 '***' 0.001 '**' 0.01 '*' 0.05 '.' 0.1 ' ' 1

Correlation of Fixed Effects:
            (Intr) stmltn
stimulatnsg -0.798
statesleep  -0.364  0.000

$emmeans
 stimulation state  emmean     SE   df lower.CL upper.CL
 tms         wake  -0.0833 0.0944 88.5   -0.271    0.104
 seeg        wake  -0.5093 0.0637 23.2   -0.641   -0.378
 tms         sleep -0.3468 0.0944 88.5   -0.534   -0.159
 seeg        sleep -0.7729 0.0637 23.2   -0.905   -0.641

Degrees-of-freedom method: kenward-roger
Confidence level used: 0.95

$contrasts
 contrast               estimate     SE    df t.ratio p.value
 tms wake - seeg wake      0.426 0.1030  40.8   4.137  0.0009 ***
 tms wake - tms sleep      0.264 0.0688 111.7   3.833  0.0012 **
 tms wake - seeg sleep     0.690 0.1238  71.9   5.569  <.0001 ***
 seeg wake - tms sleep    -0.163 0.1238  71.9  -1.313  0.5580
 seeg wake - seeg sleep    0.264 0.0688 111.7   3.833  0.0012 **
 tms sleep - seeg sleep    0.426 0.1030  40.8   4.137  0.0009 ***

Degrees-of-freedom method: kenward-roger
P value adjustment: tukey method for comparing a family of 4 estimates
```

#### Table S13: Linear Mixed Model for total HFsup (wake/sleep) (Fig 4F)
